# Oxytocin signaling regulates maternally-directed behavior during early life

**DOI:** 10.1101/2024.02.15.580483

**Authors:** Daniel Zelmanoff, Menachem Kaufman, Julien Dine, Jonas Wietek, Anna Litvin, Shaked Abraham, Savanna Cohen, Ofer Yizhar

## Abstract

Oxytocin is essential in shaping social behavior across the lifespan. While the role of oxytocin signaling in parental care has been widely investigated, little is known about its function in social behavior during early life. This is partly due to the lack of precise technologies for studying the developing brain. Here, we studied the role of oxytocin in pup social behavior under acute separation from and reunion with the mother. We show that the activity of oxytocin neurons was increased by acute maternal separation and returned to baseline after reunion. Behaviorally, maternally-separated pups emitted more ultrasonic vocalizations upon reunion, which were further modulated by nipple attachment behavior. These effects were attenuated by blocking the oxytocin receptor during maternal separation. To investigate the role of oxytocin neurons with higher precision, we established a method for transcranial optogenetic silencing of neuronal activity in untethered and freely behaving pups. Using this approach, we found that silencing of oxytocin neurons during maternal separation disrupted vocal behavior during separation and reunion in a sex-specific manner. Our findings reveal an important role of oxytocin in context-dependent vocal communication in pups, offering new insights into the mechanisms of social behavior during early life.

## Introduction

Oxytocin (OT), a nonapeptide secreted from the paraventricular (PVN) and supraoptic (SON) nuclei in the hypothalamus of mammalian brain, is widely recognized as a key neuromodulator involved in a wide array of social behaviors (reviewed in Menon and Neumann, 2023). Over the last 50 years, numerous animal studies have pointed to a central role for OT in myriad aspects of social behaviors, including pair bonding (Walum and Young, 2018), social learning (Choe et al., 2015; Dölen et al., 2013; Lewis et al., 2020), parental care (Numan and Insel, 2006; Dulac et al., 2014; Marlin et al., 2015), and adult-infant attachment (Insel and Young, 2001; Kojima and Alberts, 2011). Furthermore, animal models with disrupted OT signaling display behavioral impairments in social function, reminiscent of core symptoms in neurodevelopmental disorders such as autism spectrum disorder (ASD) (Winslow and Insel, 2002; Hörnberg et al., 2020; Peñagarikano et al., 2015; Lewis et al., 2020; Winslow et al., 2000; Takayanagi et al., 2005; Harony-Nicolas et al., 2017). While OT treatment is suggested as a potential treatment to social deficits in ASD patients (Meyer-Lindenberg et al., 2011; Young and Barrett, 2015; Insel, 2010), such clinical studies have culminated in mixed results (Sikich et al., 2021; Guastella et al., 2023; Parker et al., 2017). This suggests that our basic understanding of the OT system and its role in social behavior, particularly during early development, is incomplete.

One explanation for these inconclusive results is the notion that OT is not merely a general enhancer of pro-social behaviors but its social effects may be context-dependent. A growing list of studies have shown that apart from its pro-social effects, OT signaling can also induce aggression (Oliveira et al., 2021; Anpilov et al., 2020; Bosch et al., 2005) and social avoidance (Rogers-Carter et al., 2018; Osakada et al., 2024), and even modulate the neural and behavioral responses of social isolation (Dölen et al., 2013; Musardo et al., 2022). These findings suggest a more nuanced role for OT in social behavior, and have given rise to a modern framework of OT function, which proposes that OT enhances the salience of both positive and negative social information based on the animal’s sensory-social context (reviewed in (Shamay-Tsoory and Abu-Akel, 2016; Grinevich and Stoop, 2018; Froemke and Young, 2021; Caldwell and Albers, 2016)). During early postnatal life, an animal’s sensory-social context is unique and constantly evolving. It is characterized by close and intense interactions with parents and siblings and by increased active exploration of the surroundings, which is crucial for survival. Interestingly, this developmental stage, both in mice and in humans, is characterized by high expression of the OT receptor (OTR) in cortical and subcortical brain regions, specifically peaking in mice during the second and third weeks of life (Mitre et al., 2016; Hammock and Levitt, 2013; Newmaster et al., 2020; Rokicki et al., 2022), suggesting that OT signaling is important at this early stage. Previous studies have used chronic environmental stressors, such as maternal separation (MS) and social isolation, to test the long-term effects on the development of the OT system and social behavior (reviewed in Veenema, 2012; Sandi and Haller, 2015). Indeed, recent work has identified an important role for early life sensory experience in promoting OT-mediated changes in synaptic plasticity and behavior during development and adulthood (Zheng et al., 2014; Yu et al., 2022; Hu et al., 2022). However, very little is known about the role of OT in regulating ongoing behavioral adaptations during early life. This is partly due to a paucity of precise tools available during this sensitive period of the developing brain.

In the current study, we set out to explore the role of the OT system in mouse pups, in the context of acute separation from and reunion with the mother (also referred to as dam), as the pup-dam interaction represents a major generator of early-life sensory-social experience in mammals. While the role of OT signaling in maternal responses to such challenges is widely investigated (Marlin et al., 2015; Valtcheva et al., 2023; Schiavo et al., 2020), little is known about its involvement in homeostatic behavioral changes during early life. We acutely separated postnatal day 15-16 (hereafter referred to as P15) mouse pups from their dams and littermates for 3 hours, and studied their behavior during separation and upon reunion. By recording ultrasonic vocalizations (USVs) and nipple attachment behavior upon the reunion with an anesthetized dam, we showed that acute MS induced a robust increase in USV emission upon reunion that was further modulated by nipple attachment behavior. We found that OT neuron activity increased during acute MS and was suppressed upon reunion. Blocking the OTR during MS led to a reduction in maternally-directed behaviors in pups and altered the pattern of USV calls emitted, suggesting that OT release during MS is important for these behaviors. To investigate the role of OT release with higher spatial and temporal precision in untethered, freely behaving pups, we established an optogenetic protocol for noninvasive transcranial photoinhibition using eOPN3, a potent and highly light-sensitive opsin (Mahn et al., 2021). Using this approach, we found that optogenetic silencing of OT neurons during MS alters USV emission patterns during separation and disrupts vocal, but not non-vocal, behavior during reunion in a sex-specific manner. Our findings reveal an important role of OT in context-dependent social behavior in early life and open new avenues for mechanistic understanding of neural circuits in the pre-weaning period.

## Results

### Acute maternal separation increases maternally-directed behavior upon reunion

In dams, pup calls have been shown to induce pup-seeking behavior through the release of OT (Marlin et al., 2015; Valtcheva et al., 2023). However, the neural mechanisms engaged in the pup brain during such short-term transitions from social deprivation to engagement during early life are not well understood. To study this, we used a modified MS paradigm in which we examined reunion-related USVs and affiliative behaviors of P15 mice following three hours of MS (i.e., mother-pup assay; MP; Fig. 1A,B). In our assay, we eliminated the participation of the dam as the driving force of social interaction by studying pups’ behavior toward an anesthetized dam (Opendak et al., 2021; Zimmer et al., 2019). This procedure allowed us to identify the source of USVs and focus our investigation on the pups’ social motivation during the reunion.

**Figure 1:**
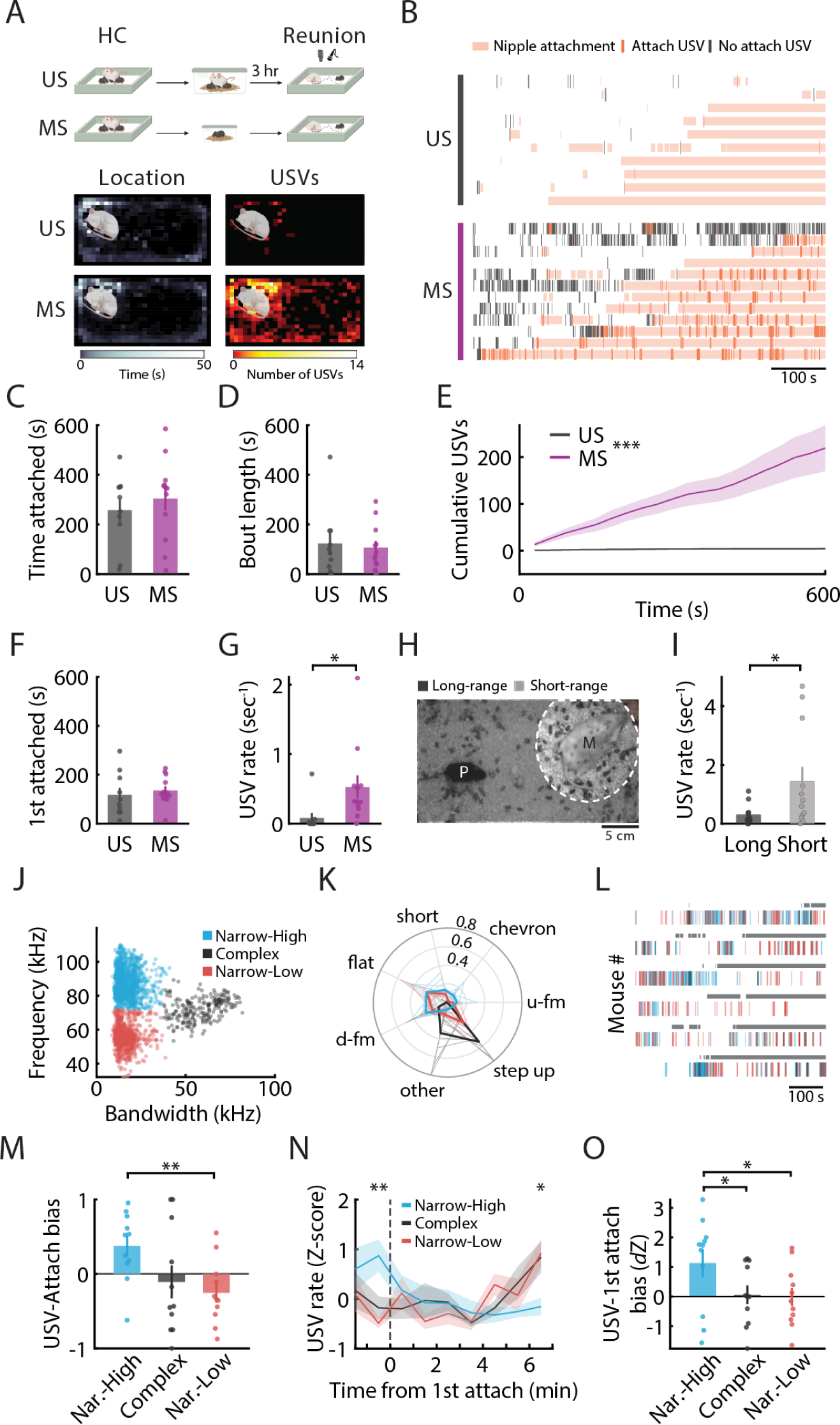
Acute maternal separation increases maternally-directed behavior upon reunion. (A) Top: Schematics of the mother-pup assay illustrating unseparated (US; n = 10) and maternally-separated (MS; n = 12) pups. Bottom: Heat maps of the location and USV counts, averaged over 1 mm^2^ bins, in relation to the dam’s position in the reunion cage. Figure 1: **(B)** Raster plot of USVs emitted upon the reunion with the dam, showing USVs emitted within (orange ticks) and outside (gray) events of nipple attachments and overlaid with bouts of nipple attachment for each pup (shaded orange). Individual mice are sorted based on total duration of nipple attachment (ascending). **(C)** Total time of nipple attachment during reunion with the dam for MS (magenta) and US (gray) pups. Student’s t-test *t*_(20)_ = *−*0.699, *P* = 0.493. **(D)** Averaged bout length of nipple attachment during reunion. Student’s t-test *t*_(20)_ = 0.331, *P* = 0.744. **(E)** Cumulative number of USVs emitted during the reunion over 30-sec bins. Mixed-design repeated-measures (RM) ANOVA. *F_groups_*_(1,20)_ = 20.814, P = 1.9 × 10*^−^*^4;^ F*_time_*_(1.216,24.327)_ = 11.375, P = 0.002; F*_groups×time_*_(1.216,24.327)_ = 10.727, *P* = 0.002. **(F)** Latency to first nipple attachment. Student’s t-test *t*_(20)_ = *−*0.57, *P* = 0.575. USV rate until the first nipple attachment. Student’s t-test *t*_(14.81)_ = *−*2.486, *P* = 0.025. A representative video still illustrating the ”mother-zone” (M) and threshold (dashed line) for long-range (dark gray) and short-range (light gray) USVs emitted by the pups (P) until the first nipple attachment. **(I)** Rate of long- and short-range USVs in MS pups until the first nipple attachment. Two-sided Wilcoxon signed-rank test *W* = 57, *P* = 0.032. **(J)** Individual USVs emitted by MS pups grouped into three clusters based on their bandwidth and mean frequency. Colors in subsequent panels correspond with this cluster analysis. **(K)** Distribution of predefined vocal categories for individual pups (thin lines) averaged across mice (bold lines) for each cluster (n*_textnormal_*_(_*_MS_*_)_ = 11). **(L)** Representative raster plots of cluster-specific USVs emitted by MS pups upon reunion. Horizontal lines (gray) represent events of nipple attachment for each pup. **(M)** The USV-attachment bias for the three USV clusters of MS pups. Higher index values indicate greater USV emission *outside* nipple attachment for the specified cluster. RM ANOVA for comparisons of different clusters with Bonferroni corrections. *F_cluster_*_(2,20)_ = 6.434, *P* = 0.007 **(N)** Normalized cluster-specific frequency of USV emission, aligned to first nipple attachment for MS pups. Two-way RM ANOVA examining the effect of cluster and time from first nipple attachment on USV emission with Bonferroni correction for post hoc comparisons. *F_cluster_*_(2,20)_ = 0.216, *P* = 0.808; *F_time_*_(7,70)_ = 2.419, *P* = 0.028; *F_cluster×time_*_(4.489,44.894)_ = 4.064, *P* = 0.005. **(O)** Quantification of cluster-specific tendency to vocalize in relation to first nipple attachment for MS pups. (RM ANOVA comparing clusters with Bonferroni-corrected post hoc comparisons: *F_cluster_*_(2,20)_ = 7.631, *P* = 0.003. Data are presented as mean *±* s.e.m. (error bars or shaded areas), \**P <* 0.05 \*\*\**P <* 0.01 \*\**P <* 0.001. For detailed statistical information, see Supplementary Table 1 and Supplementary Table 2.

An important feature of mother-pup interaction is nipple attachment, which initiates feeding interactions and is thus crucial for survival (Blass and Teicher, 1980; Jansen et al., 2008). In the reunion phase of the MP assay, pups were placed away from the anesthetized dam, and were allowed to freely explore the testing cage and engage in nipple attachment behavior. We found no differences in nipple attachment behavior in MS pups (n = 12) compared with unseparated control pups (US; n = 10; Fig. 1C,D; Extended Data Fig. 1A-D), likely due to the relatively short duration of milk deprivation in our assay (Cramer et al., 1980; Toda and Kawasaki, 2014). In contrast, we found striking differences between MS and unseparated pups in USV emission during reunion. MS pups showed robust USV emission upon reunion, primarily while in close proximity with the mother, compared with unseparated pups that displayed notably fewer USVs (Fig. 1A,B). This pattern of USV emission persisted in MS pups throughout the reunion (Fig. 1E). It was present while approaching the dam before the first nipple attachment, mainly when in close proximity to their dam (Fig. 1F-I), but also while engaging in nipple attachment (Fig. 1B). The occurrence of USV emission across varying behavioral states - both outside and during events of nipple attachment - suggests a state-dependent use of these vocalizations (Sangiamo et al., 2020). We, therefore, examined next whether these distinct behavioral states might be associated with unique types of USVs.

We used machine-learning-based tools to study the USV emission patterns recorded from MS pups during the reunion phase (Fonseca et al., 2021; Coffey et al., 2019; Extended Data Fig. 1E-H; Extended Data Fig. 2). We identified three profiles of USVs emitted by MS pups: 1) USVs with narrow bandwidth and high mean frequency (”Narrow-High”), 2) USVs with varied bandwidth and medium mean frequency (”Complex”) and, 3) USVs with narrow bandwidth and low mean frequency (”Narrow-Low”; Fig. 1J). In addition, we compared our classification method with a commonly used approach that assigns USVs to predefined vocal categories based on their spectro-temporal properties (Fonseca et al., 2021; Grimsley et al., 2011). This revealed a considerable overlap of some vocal categories, mainly in the Narrow-High and Narrow-Low USVs, and yet significant differences in the distribution of categories across the three USV profiles (Fig. 1K; Extended Data Fig. 1I-J), suggesting that each profile represents a unique set of USV subtypes within the pup’s vocal repertoire.

We next asked whether these USV profiles are differentially linked to distinct nipple attachment states. For this purpose, we calculated the USV-attachment bias for each cluster, indicating the tendency of MS pups to use a specific USV cluster outside or during an attachment event. This analysis revealed significant differences in the dynamics of USV clusters during attachment behavior. We found that MS pups emitted more ”Narrow-High” USVs *outside* of attachment events, compared with ”Narrow-Low” USVs that were more common *during* attachment events (Fig. 1L-M). Specifically, ”Narrow-High” USVs were more prevalent before the first attachment event and gradually declined during the trial (Fig. 1N-O). In contrast, ”Narrow-Low” and ”Complex” USVs were less common before the first attachment event and showed a slight increase toward the end of the trial. Overall, we found that acute MS induces robust USV emission upon reunion with the dam, which is further modulated by nipple attachment behavior.

### OT neuron activity is increased by MS and suppressed upon reunion

We next set out to explore the role of the OT system in MS-induced maternally-directed behavior. The OT system is known to be involved in the regulation of social interactions that essentially require close contact with conspecifics, such as social recognition, maternal care, pair bonding and aggression (Kojima and Alberts, 2011; Oliveira et al., 2021; Dölen et al., 2013; Oettl et al., 2016; Tang et al., 2020b; Insel, 2010; Anpilov et al., 2020; Marlin et al., 2015). However, the involvement of the OT system in the acute experience of social deficit, particularly during early life, remained less explored. Pioneering work by James Winslow and Thomas Insel has laid the foundation for understanding OT’s role in MS-induced USVs in early life (Winslow et al., 2000; Winslow and Insel, 2002; Insel and Winslow, 1991). Recently, OT was shown to play a conserved role in affiliative behavior toward the dam in both rodents and monkeys, although not in USV emission during MS (Liu et al., 2023b). Further research showed that OT signaling drives the changes in social behavior following one week of post-weaning social isolation in mice (Musardo et al., 2022). Altogether, these findings suggest a potential role for the OT system in social transitions, specifically the shift from separation to reunion. We therefore hypothesized that changes in the pup’s motivational drive to seek and maintain social interaction with the dam are linked with the activation of OT neurons during MS.

To address this question, we assessed the neuronal activation of the hypothalamic OT-expressing cells in the context of MS. We labeled the transcription factor c-Fos, an indirect marker for neuron activity, in PVN and SON sections from P15 pups after two hours of MS (n_(MS)_ = 5), compared to unseparated pups (n_(US)_ = 4), and analyzed the selective activation of OT neurons by colocalization of OT and c-Fos immunoreactive cells (Fig. 2A). Our results showed increased activity of OT neurons both in the PVN and SON of MS pups compared with unseparated pups (Fig. 2B). This elevated activity was attenuated in pups that underwent 3 hours of MS followed by 1.5 hours of reunion in their home-cage (”Re”; n_(Re)_ = 4). Other brain regions that are known to be connected with the PVN and involved in stress response, namely the paraventricular thalamus (PVT), lateral habenula (LHab) and suprachiasmatic nucleus (SCN), did not show elevated activity in MS pups compared with control, suggesting these changes were not due to a global brain activation (Fig. 2C).

**Figure 2:**
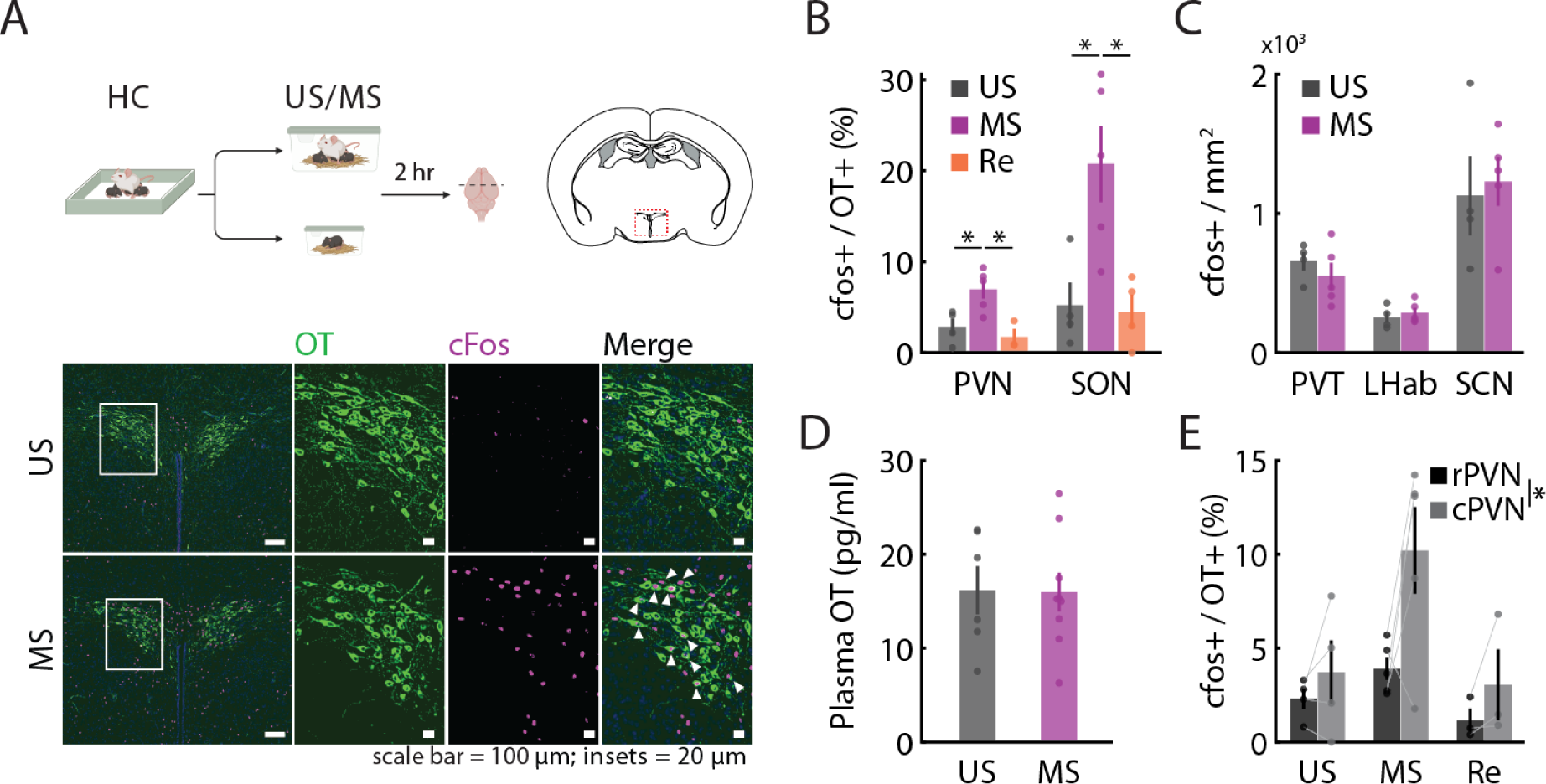
OT neuron activity is increased by maternal separation and suppressed upon reunion. **(A)** Top: Schematics of the experimental design (MS pups: n_(MS)_ = 5; US pups: n_(US)_ = 4). Red dash indicates the location of the PVN (top right). Bottom: Representative images of the PVN from US and MS pups. White arrowhead indicates colocalization of OT (green) and c-Fos (magenta). **(B)** OT neurons positive for c-Fos immunoreactivity in the PVN and SON of US (gray), MS (magenta) and reunited pups (Re pups; orange; n_(Re)_ = 4). One-way ANOVA with Bonferroni corrected post hoc comparisons. PVN: *F_groups_*_(2,9)_ = 7.981, *P* = 0.01; SON: *F_groups_*_(2,10)_ = 8.318, *P* = 0.007. **(C)** Quantification of c-Fos positive cells (normalized to area) in the paraventricular thalamus (PVT), lateral habenula (LHab) and suprachiasmatic nucleus (SCN) for US and MS pups. Student’s t-test: *t_PV T_* _(7)_ = 0.868, *P* = 0.414; *t_LHab_*_(7)_ = *−*0.619, *P* = 0.555; *t_SCN_*_(7)_ = *−*0.315, *P* = 0.762. **(D)** Comparison of plasma OT concentration between US and MS pups. Student’s t-test *t*_(13)_ = 0.063, *P* = 0.951. **(E)** Quantification of c-Fos positive OT neurons in the rostral PVN (rPVN; [*−*0.58: *−*0.94]*mm* relative to bregma) and the caudal PVN (cPVN; [*−*1.06: *−*1.22] *mm* relative to bregma) for US and MS pups. Mixed-design RM ANOVA *F_rostal/caudal_*_(1,9)_ = 6.912, *P* = 0.027; *F_groups_*_(2,9)_ = 5.305, *P* = 0.03; *F_groups×_*_[_*_rostal/caudal_*_](2,9)_ = 1.890, *P* = 0.206. Data are presented as mean *±* s.e.m. (error bars), \**P <* 0.05. For detailed statistical information, see Supplementary Table 2.

The elevated activity of OT neurons observed in MS pups was not associated with an increase in peripheral OT concentration (Fig. 2D). We considered the possibility that OT neurons are not yet mature enough to release OT into the circulation at this age. However, intraperitoneal (i.p.) injection of the blood brain barrier-impermeable retrograde tracer Fluoro-Gold (FG; Merchenthaler, 1991) in P7 pups revealed selective labeling of magno-OT neurons (FG+/OT+; Extended Data Fig. 3A) and distribution of parvo-OT neurons mainly in the caudal PVN (FG-/OT+), consistent with their distribution in adult mice (Lewis et al., 2020). Consistent with the lack of elevation in systemic OT, we found that activated OT neurons were mainly distributed in the caudal PVN (Fig. 2E).

Previous work has demonstrated synaptic changes in PVN neurons of adult mice following a brief exposure to a stressor (Sterley et al., 2018). We therefore asked whether PVN-OT neurons show changes in their intrinsic or synaptic properties following MS. We were able to classify the PVN-OT population of pups into magno- and parvo-OT subtypes based on their established electrophysiological signatures (Luther and Tasker, 2000; Lewis et al., 2020; Extended Data Fig. 3B-D), suggesting that OT neurons show adult-like electrophysiological features as early as the second week of life. However, we found no differences in synaptic input as well as intrinsic properties of OT neurons in MS pups compared with unseparated pups (Extended Data Fig. 3E-G; see Methods), suggesting that the behavioral changes associated with MS are not caused by changes in synaptic input or altered excitability of OT neurons.

Altogether, we show that MS increases the activity of hypothalamic OT neurons that is further attenuated upon reunion. Specifically, our results demonstrate that the magno-OT and parvo-OT neurons are anatomically and functionally mature as early as the second week of life in mice, and suggest a selective role for parvo-OT neurons in MS.

### Blocking OT receptors during MS inhibits maternally-directed behavior upon reunion

Having observed increased activity of OT neurons during MS, we next hypothesized that blocking the OTR during MS would alter maternally-directed behavior. We injected P15 pups with an OTR antagonist (OTRA; 10mg/kg; 10 ul/g i.p.) 30 minutes before MS (“OTRA”; n_(OTRA)_ = 21) and assessed their maternally-directed behavior as compared to vehicle-injected control pups (“SL”; 95% normal saline, 5% DMSO; n_(SL)_ = 21; Fig. 3A). In pups treated with OTRA we found a profound decrease in the total duration of nipple attachment and a delayed phenotype of attachment events compared with control pups (Fig. 3B). These differences could not be explained by changes in event length distribution, attachment event intervals, or the total number of events in OTRA pups (Extended Data Fig. 4A-C).

**Figure 3:**
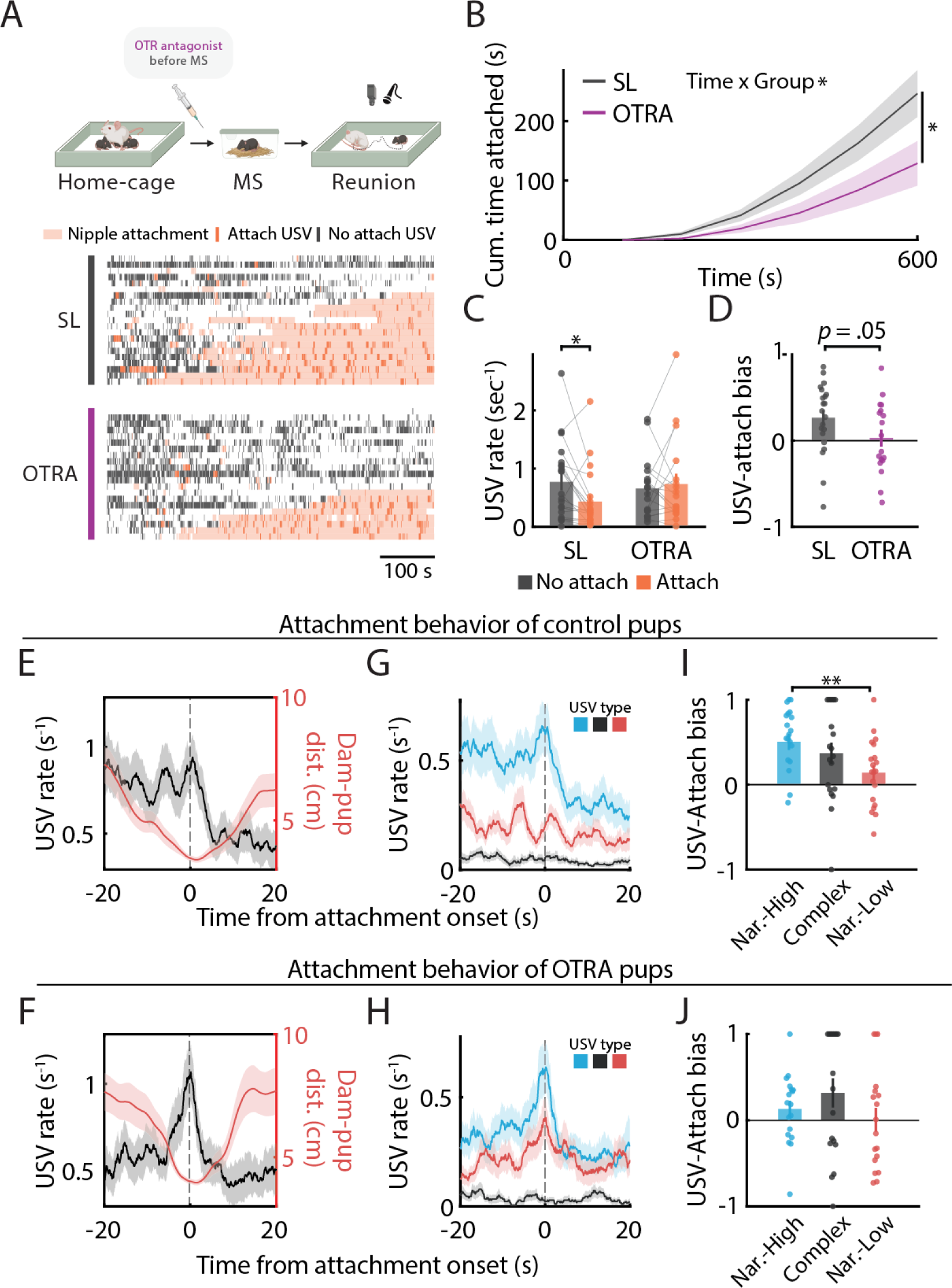
Blocking the OT receptor during maternal separation suppresses maternally-directed behavior upon reunion. Figure 3: **(A)** Schematic representation of the experimental protocol. P15 pups were treated with OTRA (n_(OTRA)_ = 21; i.p.) and compared to vehicle-treated mice (SL; n_(SL)_ = 21). Bottom: Raster plot depicting USVs emitted by OTRA-treated (magenta) and control (gray) pups, overlaid with bouts of nipple attachment upon reunion with the dam and sorted based on total duration of nipple attachment (ascending). **(B)** Cumulative time of nipple attachment. Mixed-design repeated measures ANOVA *F_time_*_(1.12,44.811)_ = 42.786, *P* = 1.75 *×* 10^-8^; *F_groups_*_(1,40)_ = 4.574, *P* = 0.039; *F_time×groups_*_(1.12,44.811)_ = 4.068, *P* = 0.045. **(C)** Quantification of USV rate outside (gray) and within (orange) attachment events. Mixed-design repeated measures ANOVA with post hoc Bonferroni correction *F_attach_*_(1,37)_ = 1.842, *P* = 0.183; *F_groups_*_(1,37)_ = 0.299, *P* = 0.588; *F_attach×groups_*_(1,37)_ = 4.565, *P* = 0.039. **(D)** The USV-attachment bias. Two-sided Mann Whitney U-test *U* = 119, *P* = 0.05. **(E–H)** Temporal dynamics of USV emission around the onset of attachment events for control (n = 84 events in 21 mice; E,G) and OTRA-treated pups (n = 59 events in 18 mice; F,H). **(E–F)** Peri-event time histogram (PETH) of average USV rate (black) and mother-pup distance (red) centered at the onset of nipple attachment. **(G–H)** PETH of averaged USV rate of the three USV clusters (as shown in Fig.5) centered at the onset of nipple attachment. Blue trace: narrow bw, high frequency (”Nar.-High”); black trace: mixed bw, medium frequency (”Complex”); red trace: narrow bw, low frequency (”Nar.-Low”). **(I– J)** Cluster-specific USV-attachment bias for control (I) and OTRA (J) pups. RM ANOVA for comparisons of different clusters with Bonferroni corrections *F_SL_*_(2,38)_ = 4.669, *P* = 0.015; *F_OT RA_*_(2,38)_ = 1.624, *P* = 0.212. Data are presented as mean *±* s.e.m. (error bars or shaded areas), \**P <* 0.05, \*\**P <* 0.01. For detailed statistical information, see Supplementary Table 2.

Next, we examined the USVs emitted upon reunion with the dam. Our results revealed no alterations in the number of USVs between OTRA-treated and control pups (Extended Data Fig. 4D). OTRA-injected pups emitted a rich repertoire of USVs comparable to control pups (Extended Data Fig. 5A,B,E,F). However, we found significant changes in the physical properties of the vocalizations (Extended Data Fig. 4D). Specifically, we found that OTRA pups emitted USVs with higher mean frequency and lower amplitude compared with control, with no differences in duration or bandwidth (Extended Data Fig. 4E-H). These findings indicate that inhibition of OT signaling can induce robust changes in nipple attachment and in the properties of emitted USVs. This is consistent with accumulating evidence demonstrating that USV emission can be modulated by emotional and social states (Chen et al., 2021a; Sangiamo et al., 2020; Knutson et al., 2002; Panksepp, 2005).

To understand the effect of OTR inhibition on the relationship between USV emission and nipple attachment, we first evaluated the USV rate of pups outside and within attachment events. Control pups displayed a lower rate of USV calls during nipple attachment than outside of attachment events. In contrast, OTRA-treated pups emitted similar numbers of USVs in these two states (Fig. 3C-D). We next examined the temporal dynamics of USV emission around the onset of nipple attachment events. In both OTRA-treated and control pups, nipple attachment was preceded by a shortening of dam-pup distance (Fig. 3E,F). Notably, in control pups this decrease in dam-pup distance was associated with high USV emission rates, which were rapidly attenuated upon attachment. However, OTRA-treated pups displayed a different USV emission pattern, characterized by a lower USV rate before attachment, an abrupt peak around attachment onset, followed by a decline to pre-attachment USV rates. Using the classification method described in Fig. 1J, we found that this attachment-associated modulation of USVs was mainly driven by the Narrow-High subset of USVs (Fig. 3G,H). Quantification of this phenotype revealed that control pups emitted Narrow-High USVs predominantly outside attachment events compared with the Narrow-Low USVs, while in OTRA-treated pups this bias was abolished (Fig. 3I,J).

We also examined more closely USV modulation around the first event of nipple attachment in MS pups after treatment with OTRA. Although treatment with OTRA did not induce a significant delay in first nipple attachment, it did alter the dynamics of USV emission while approaching the dam prior to the first nipple attachment. Specifically, while control pups increased their USV emission rate when approaching the dam, OTRA pups did not show such modulation (Extended Data Fig. 4I-J). We further found that control pups frequently used Narrow-High USVs before the first attachment, and this USV class rapidly declined after attachment. In contrast,the Narrow-Low and Complex USVs were significantly less modulated by the first attachment (Extended Data Fig. 5C-D). This cluster-specific modulation of USVs by the first nipple attachment was absent in OTRA-treated pups (Extended Data Fig. 5G-H). Taken together, these findings suggest that activation of the OT system and OT signaling during acute MS are essential for the manifestation of maternally-directed behavior in early life.

### Development of a method for transcranial optical silencing of OT neurons in postnatal mice

The activation of OT neurons during MS, together with OTRA-induced changes in maternally-directed behavior during reunion, led us to hypothesize that activation of the OT system and central release of OT, specifically during acute MS, are essential for the manifestation of maternally-directed behavior upon reunion. However, the pharmacokinetics of OTRA treatment did not allow us to dissociate the specific role of OT release during MS from a potential role during reunion. To address this question with higher temporal precision, we aimed to develop an optogenetic approach for neuronal silencing in the pup brain. However, the conventional optogenetic approach, which requires bulky implants and fiber optic tethering, is very challenging to achieve and highly disruptive in pre-weaning mice that are co-housed with their dam and littermates. We therefore set out to establish an approach for non-invasive, transcranial, tether-free optogenetics in pre-weaning mice.

The main challenge in transcranial optogenetics is the limited light penetrance to deep brain structures, mainly at the blue-green wavelength range, due to increased scattering and absorption in skin, skull and neural tissue (Tromberg et al., 2000). Inhibitory optogenetic tools are particularly challenging to apply transcranially, since they typically require continuous light delivery and possess an intrinsically low light sensitivity (Wiegert et al., 2017). We hypothesized that eOPN3, a red-shifted and light-sensitive opsin recently developed by our group (Mahn et al., 2021), could alleviate many of these constrains. We first confirmed, using autaptic neuron recordings, that illumination with low-irradiance red light (636 nm) was sufficient to evoke robust reduction in postsynaptic currents (PSCs) in eOPN3-expressing hippocampal neurons *in vitro* (Fig. 4A-C). These results indicated that eOPN3 may serve as an efficient tool for transcranial inhibition, particularly for prolonged silencing as required in MS.

**Figure 4:**
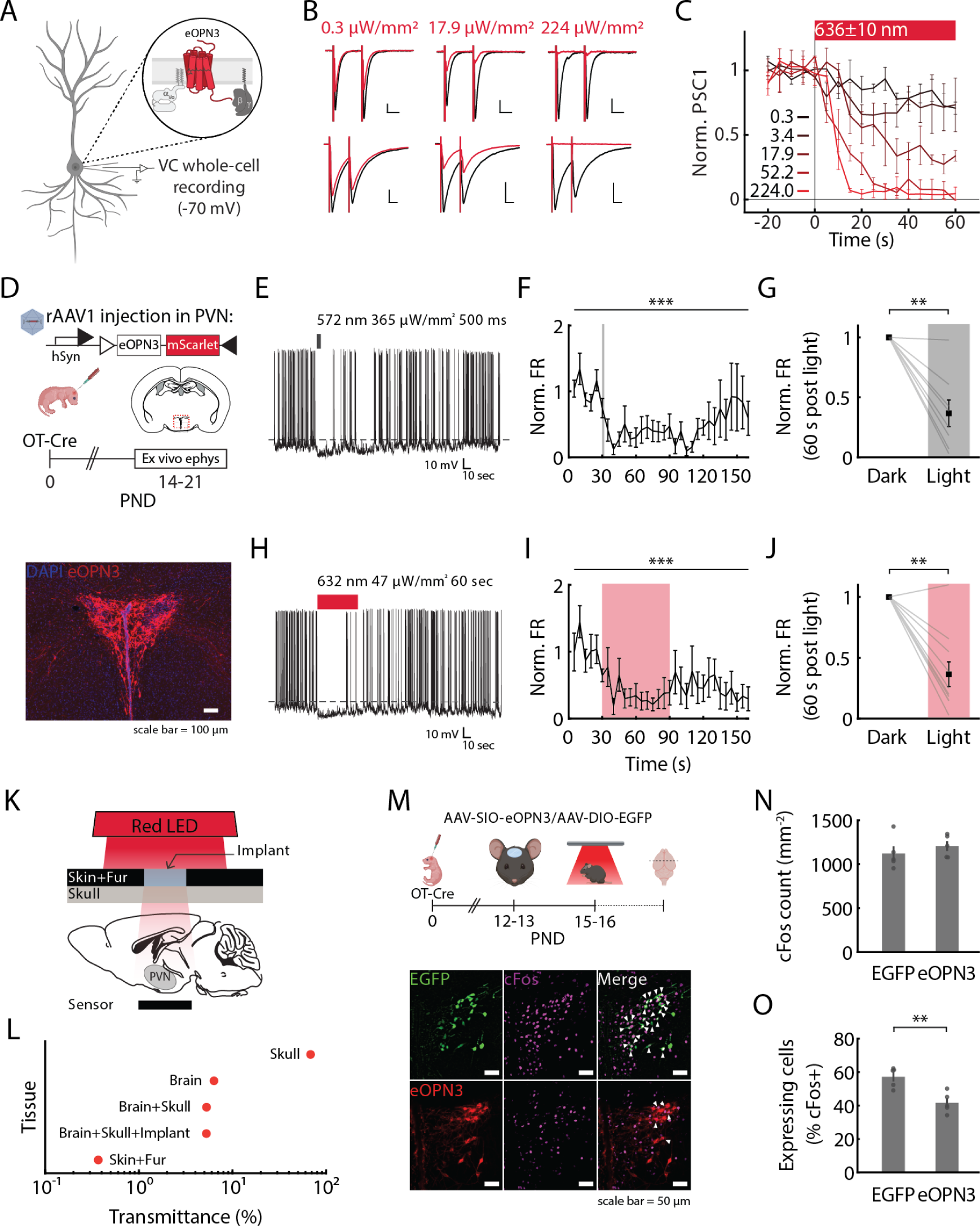
eOPN3-mediated transcranial photoinhibition of OT neurons in freely behaving pups. **(A)** Schematic illustrating the electrophysiological recordings in autaptic hippocampal neurons expressing the G_i/o_ protein-coupled opsin eOPN3. Figure 4: **(B)** Representative whole-cell voltage clamp recordings from glutamatergic (top) and GABAergic (bottom) neurons of responses to pairs of depolarizing current injections in the dark (black) or during red light illumination (636 *±* 10 *nm*) at three intensities. **(C)** Summary of time course over 5-sec bins, showing postsynaptic currents (PSCs) before and during 60 seconds of illumination at the indicated red light intensities (EC_50_ = 4.6 µW/mm^2^). Values represent light irradiance in µW/mm^2^. **(D)** Top: Schematic of neonatal viral injection and electrophysiological recordings from acute slices of the PVN (dashed line) containing OT-eOPN3 neurons. Bottom: Representative image showing early (i.e., P14) somatic and axonal expression of eOPN3 opsin in PVN-OT neurons. **(E–J)** Representative traces (E and H), summary of time course over 5-sec bins (F and I), and normalized post-illumination comparison (G and J) of firing rate in neurons expressing OT-eOPN3 before and after continuous stimulation with green (E-G; n = 8) or red (H-J; n = 10) light, respectively. (F and I) Friedman test for comparison of firing rate over time. 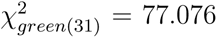 (G and J) Paired t-test *t_green_*_(7)_ = 5.273, *P* = 0.001; *t_red_*_(9)_ = 4.053, *P* = 0.003. **(K)** Diagram illustrating the characterization of red light penetration through brain tissue. **(L)** Transmittance of red light through the different tissues. **(M)** Top: Schematic representation of the experimental approach for eOPN3-mediated transcranial silencing of OT neurons in freely moving pups. Bottom: representative images of the PVN from OT-EGFP (green) and OT-eOPN3 (red) mice co-stained for c-Fos (magenta) after salt loading. Arrowheads indicate the overlap of c-Fos-positive neurons and EGFP-or eOPN3-positive neurons. **(N)** Quantification of c-Fos positive cells in the PVN. Student’s t-test *t*_(8)_ = *−*0.847, *P* = 0.421. **(O)** Percentage of EGFP-and eOPN3-expressing neurons positive for c-Fos immunoreactivity in the PVN. Student’s t-test *t*_(8)_ = 4.034, *P* = 0.004. Data are presented as mean *±* s.e.m. (error bars), \*\**P <* 0.01 \*\*\**P <* 0.001.

Next, we examined whether optogenetic silencing with eOPN3 could effectively inhibit neuron activity in PVN-OT neurons in early life. To achieve early life cell-type-specific expression of eOPN3 in PVN-OT neurons, we injected a Cre-dependent viral vector (AAV-hSyn-SIO-eOPN3-mScarlet) to P0 pups from a mouse strain that endogenously expresses Cre recombinase under the control of OT promoter (OT-Cre; Fig. 4D, top). We confirmed the expression of eOPN3 in the PVN-OT neurons (OT-eOPN3) two weeks after injection (Fig. 4D, bottom). To establish the functional expression of eOPN3 in OT neurons, we performed whole-cell recordings from eOPN3-expressing OT neurons in acute PVN slices. As expected, illumination of eOPN3-expressing cells with green light (572 nm, 365 µW/mm^2^ for 500 ms; n = 8 cells) evoked a potent and reversible reduction in firing rate (Fig. 4E-G). Consistent with our hippocampal neuron recordings, activation of OT-eOPN3 neurons with red light (632 nm) at very low irradiance (47 µW/mm^2^ for 60 sec) was also sufficient to evoke a significant reduction in action potential firing (n = 10 cells; Fig. 4H-J). We concluded that delivery of low irradiance red light is sufficient to stimulate the highly sensitive eOPN3 opsin and suppress OT neural activity.

Next, we determined whether we could deliver a sufficient amount of red light to the PVN, located 4-4.5 mm below the skull surface in P15 mice, via a minimally-invasive optogenetic strategy. To test this, we first designed a minimally-invasive surgical procedure that allowed stable optical access to the pups’ intact skull and optimized the amount of light that could reach the PVN (Extended Data Fig. 6A). Pups with an intact-skull implant showed minimal damage to the skull and brain tissue, displaying high rates of successful recovery from the surgical procedure and long-term survival when monitored up to weaning.

To measure the amount of red light that can reach the PVN, implanted pups were euthanized at P15 and decapitated for *ex vivo* measurements (Fig. 4K). A custom red LED array was installed at a distance of 20 cm above the testing surface to detect the amount of red light transmitted through the entire brain (see Methods; Extended Data Fig. 6B). Our measurements revealed that light transmission through the implant, skull and brain (transmittance = 5.25%) was similar to transmission through the brain and skull or the extracted brain only (5.28% and 6.33%, respectively; Fig. 4L). These values were in line with the low transmission of red light through the mouse skin and fur (0.37%) and the high transmission through the isolated skull (68.1%), supporting our surgical approach. Furthermore, with this illumination setup we detected an absolute irradiance of 8.77 µW/mm^2^ below the ventral surface of the brain in implanted mice. In light of our *in vitro* and *ex vivo* experiments, this indicates that a sufficient amount of light reaches the PVN to activate eOPN3 under these conditions.

### eOPN3 facilitates transcranial, tether-free photoinhibition of OT neurons in freely-behaving pups

Next, we investigated whether the transcranial optogenetic strategy described above could be used to inhibit PVN-OT neurons in pre-weaning pups *in vivo*. To obtain a high-activity baseline in OT neurons, we used a salt loading procedure, previously demonstrated to evoke robust c-Fos expression in OT neurons (Hoffman et al., 1993; Hasan et al., 2019). As expected, P15 pups systemically injected with hypertonic solution (i.e., salt-loading; NaCl 9%; i.p.) displayed increased c-Fos immunoreactivity in both OT-positive and OT-negative neurons in the PVN and SON, compared with control pups injected with normal saline (NaCl 0.9%; Extended Data Fig. 7A).

We expressed eOPN3 in PVN-OT neurons by injecting AAV-hSyn-SIO-eOPN3-mScarlet into the PVN of OT-Cre P0 mice (OT-eOPN3; n = 5; Fig. 4M). Control OT-Cre pups were injected with AAV-hSyn-DIO-EGFP into the PVN (OT-EGFP; n = 5). We then asked whether transcranial illumination (655 nm; continuous for 90 minutes) could suppress c-Fos immunoreactivity in PVN-OT neurons of salt-loaded pups. Strikingly, while OT-eOPN3 pups showed no change in the general c-Fos count in the PVN compared with OT-EGFP controls, we observed a significant suppression of c-Fos immunoreactivity in PVN OT-eOPN3-expressing neurons compared with controls (Fig. 4N-O).

We further reaffirmed these findings in a separate cohort, in which OT-eOPN3 (n = 4) and OT-EGFP (n = 5) pups were anesthetized at P15 and underwent continuous illumination via a fiber-coupled red LED (630 nm; 150 mW/mm^2^) placed above the implant (Extended Data Fig. 7B). While overall PVN activity remained unchanged, we found that red light illumination decreased the activity of PVN-OT neurons in OT-eOPN3 pups compared to OT-EGFP pups (Extended Data Fig. 7C-D). Importantly, the expression of eOPN3 in itself did not cause a change in c-Fos counts in the absence of red light illumination, suggesting that this effect was not due to constitutive activity of eOPN3 (Extended Data Fig. 7D). Consistent with previous work (Cardin et al., 2010), we did not observe any evidence of overheating or photodamage to the brain tissue under these illumination conditions. Taken together, these data demonstrate that eOPN3 offers a reliable and safe method for transcranial photoinhibition of OT neurons in untethered freely-behaving pups.

### Optogenetic silencing of OT neurons during MS selectively alters vocal behavior during MS and reunion

Next, we sought to assess whether transcranial inhibition of OT neurons could modulate pups behavior. We bilaterally expressed eOPN3 (n = 19) or EGFP (n = 26) in hypothalamic OT neurons of P15 OT-Cre pups of both sexes (OT-EGFP: n_(F)_ = 15, n_(M)_ = 11; OT-eOPN3: n_(F)_ = 9, n_(M)_ = 10), and used the transcranial approach described above to suppress OT neuron activity during 3 hours of MS (see Methods). We then examined the behavior of pups during separation and reunion with the dam (Fig. 5A). We observed unilateral or bilateral expression of eOPN3 also in the SON of 15 out of 19 pups (Extended Data Fig. 8A), indicating that our manipulation adequately covers all of the OT neurons shown to be activated during MS.

**Figure 5:**
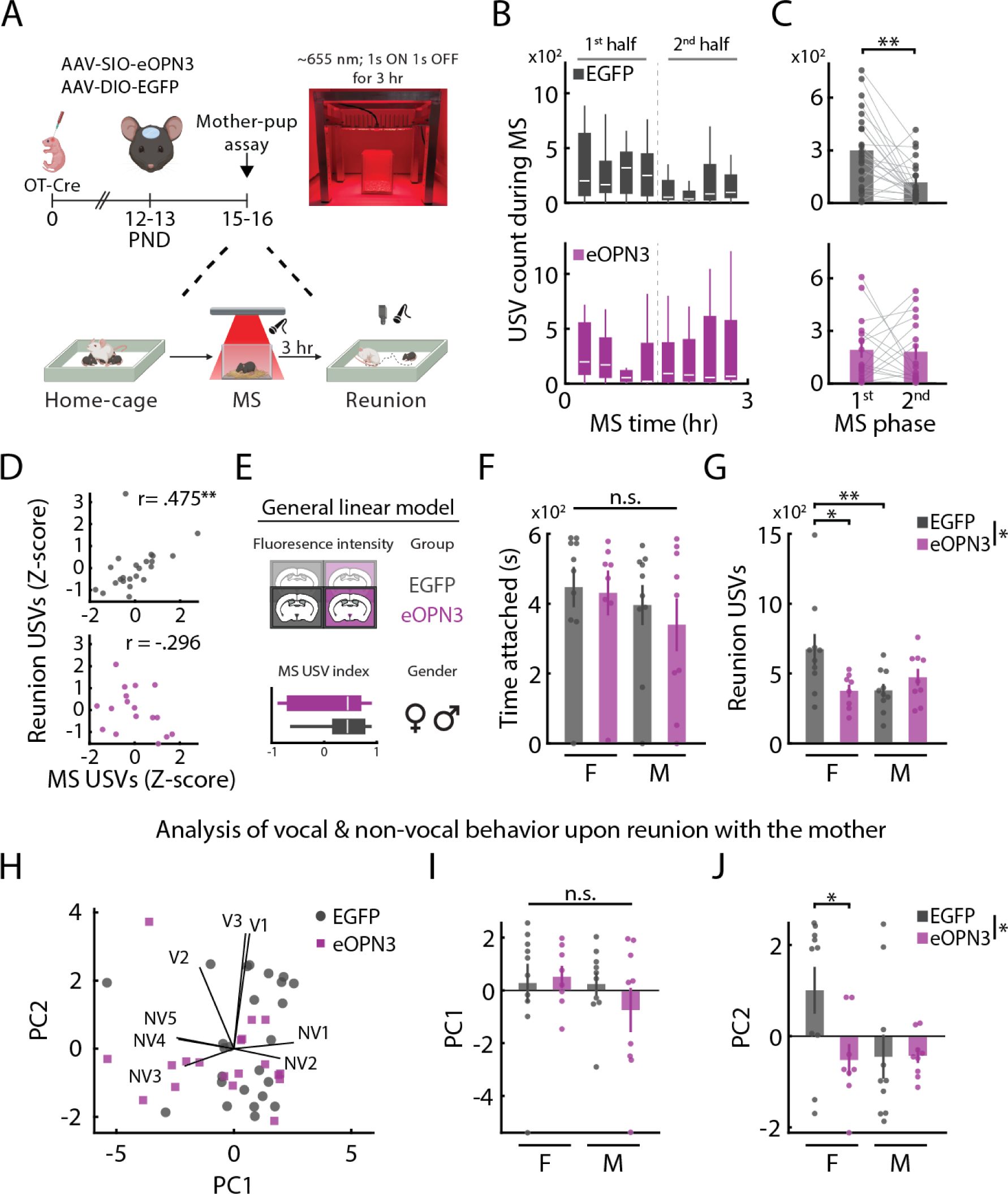
Optogenetic silencing of OT neurons during MS selectively alters vocal behavior of pups during separation and reunion. **(A)** Schematic representation of the experimental approach, as described in Fig. 4. P15 pups underwent optogenetic silencing of OT neurons during MS (n_(OT-EGFP)_ = 26, n_(OT-eOPN3)_ = 19). Image shows the red LED illumination setup. Figure 5: **(B–C)** Analysis of USV emission during MS for OT-EGFP (gray; n = 24) and OT-eOPN3 (magenta; n = 17) pups. (B) USV counts during a 3-hour MS period, recorded in 5-min segments at 15-min intervals. Box plots mark IQR (box edges) and median (white line); whiskers mark 1.5*±* IQR. Mixed-design repeated measures ANOVA *F_group_*_(1,37)_ = 0.012, *P* = 0.912; *F_sex_*_(1,37)_ = 0.513, *P* = 0.478. (C) Summary of USV count during the first and second halves of MS. Mixed-design repeated measures ANOVA with post-hoc Bonferroni correction comparing MS phases *F_phase_*_(1,37)_ = 5.955, *P* = 0.02; *F_group_*_(1,37)_ = 0.245, *P* = 0.624; *F_group×phase_*_(1,37)_ = 4.743, *P* = 0.036. **(D)** Correlation of USV counts during MS and reunion for OT-EGFP (top; Kendall tau coefficient; *P* = 0.002) and OT-eOPN3 (bottom; *P* = 0.108) pups. **(E)** Schematic outline of the general linear model (GLM) used for predicting maternally-directed behavior upon reunion for OT-EGFP (n_(F)_ = 10, n_(M)_ = 10) and OT-eOPN3 (n_(F)_ = 8, n_(M)_ = 9) pups. The following data are categorized by gender. **(F)** Total time of nipple attachment during reunion with the dam. GLM fitting: 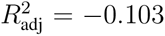; *P_group_* = 0.873; *P_sex_* = 0.599; *P_group×sex_* = 0.759. **(G)** USV counts during reunion. GLM fitting with post-hoc comparisons of Bonferroni-corrected estimated marginal means: 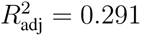; *P_group_* = 0.012; *P_sex_* = 0.002; *P_group×sex_* = 0.005. **(H)** Principal component analysis (PCA) of maternally-directed behaviors upon reunion for OT-EGFP and OT-eOPN3 pups. Each data point represents an individual pup, and arrows indicate the projection of non-vocal (NV1-NV5) and vocal (V1-V3) behaviors onto PC1 and PC2. **(I–J)** The distribution of PC1 (I) and PC2 (J) fitted to the model. GLM fitting with post-hoc comparisons of Bonferroni-corrected estimated marginal means. For PC1: 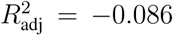; *P_group_* = 0.834; *P_sex_* = 0.985; *P_group×sex_* = 0.357; for PC2: 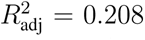; *P_group_* = 0.024; *P_sex_* = 0.007; *P_group×sex_* = 0.044. Data are presented as mean *±* s.e.m. (error bars), \*\**P <* 0.01 \*\*\**P <* 0.001. For detailed statistical information, see Supplementary Table 2.

First, we asked whether OT neuron inhibition during MS would modulate USV emission during separation (MS-USVs). We found that, while USV emission spontaneously decreased in OT-EGFP pups over the course of the 3-hour MS period, dropping significantly during the second half of this period (Fig. 5B-C, top), OT-eOPN3 pups continued to vocalize at a similar rate, with no observed difference in USV counts between the first and second half of MS (Fig. 5B-C, bottom). When accounting for sex, our analysis did not reveal a significant difference in USV counts between male and female pups, nor was there interaction with sex (Supplementary Table 2). Notably, the overall MS-USV counts for OT-eOPN3 pups were not significantly different from the control pups, indicating that activation of the OT system during MS primarily affects the dynamics of MS-USVs, rather than the total count of USVs, and suggesting a modulatory effect of OT on USV emission during MS.

Based on our previous findings, we then hypothesized that activation of the OT system during MS plays an important role in guiding maternally-directed behavior upon reunion. We therefore tested whether optogenetic silencing of OT neurons during MS would alter this behavior. Analysis of USV counts during reunion (Reunion-USVs) revealed that, while control pups exhibited a strong positive correlation between MS-USVs and Reunion-USVs, this correlation was absent in pups expressing eOPN3 (Fig. 5D).

We then leveraged the rich dataset we collected in the current experimental design, and evaluated the extent to which certain features contributed to the variation in behavior upon reunion. Specifically, in addition to the categorical information about the experimental group (i.e., OT-eOPN3 vs. OT-EGFP) and sex, we analyzed the USV attenuation index during MS and the viral expression levels in the mouse PVN (Extended Data Fig. 8B-C). To determine the influence of pups’ categorical, behavioral and histological features described above on the manifestation of maternally-directed behavior during reunion, we used mice with comprehensive histological and behavioral data from MS and reunion, and fitted a general linear model to predict the behavior (n = 37; Fig. 5E; see Methods). This analysis revealed that none of the predictive features could account for the variance in nipple attachment behavior (Fig. 5F). In contrast, both the experimental group and sex independently predicted Reunion-USVs. Remarkably, female OT-eOPN3 pups displayed significant suppression in Reunion-USVs when compared to female OT-EGFP pups, whereas male OT-eOPN3 pups emitted USVs similar to their control counterparts. Furthermore, while we observed a marked difference in Reunion-USVs between female and male EGFP-OT pups, this sexual dimorphism was absent in OT-eOPN3 pups (Fig. 5G).

We then categorized measurements of pup behavior toward the dam into non-vocal (”NV”) and vocal (”V”) behaviors, finding strong correlation within each domain (Extended Data Fig. 9A). Therefore, we employed principal component analysis (PCA) to project behavioral variables recorded during reunion onto the first two principal components (PCs) that explained approximately 77% of the variance (Fig. 5H, Extended Data Fig. 9B). Notably, non-vocal and vocal behaviors distinctly projected onto PC1 and PC2, respectively, highlighting their distinct nature in this context. Consistent with our previous results, we found that, while none of the predictive features could account for the variance in PC1 (Fig. 5I), the variance in PC2 was significantly predicted by both the experimental group and sex. Specifically, female OT-eOPN3 pups, but not male OT-eOPN3 pups, exhibited significant alteration in PC2 when compared to their control counterparts (Fig. 5J). Taken together, our findings underscore the critical role of OT neuron activity during MS in shaping vocal, but not non-vocal, behavior during separation and subsequent reunion with the dam. To the best of our knowledge, these results comprise the first demonstration of a sex-specific contribution of OT neurons to maternally-directed behavior in the pre-weaning period.

## Discussion

Our current understanding of the OT system’s role in social behavior is derived mostly from studies conducted in the adult brain, in which sensory and social networks are mature (Menon and Neumann, 2023; Grinevich and Stoop, 2018; Froemke and Young, 2021). Despite the prominent expression of OT receptors during early life (Hammock and Levitt, 2013; Mitre et al., 2016; Newmaster et al., 2020; Rokicki et al., 2022), and the crucial role of social experience during this sensitive period (Veenema, 2012; Sandi and Haller, 2015), very little is known regarding the ontogeny of the OT system and its involvement in the process of shaping early-life social behavior (Grinevich et al., 2014). This gap is largely due to limited knowledge of the full capacity of social behavior in the developing postnatal brain, and the scarcity of suitable technologies for circuit-based intervention during this critical period. In the current study, we investigated the role of OT signaling in context-dependent early-life social behavior, combining detailed behavioral analysis, pharmacological interventions, and a novel optogenetic approach for transcranial silencing in unthetered freely behaving pups.

Neonatal USVs in rodents, generally viewed as distress calls, have traditionally been investigated in the controlled context of social isolation (Elwood and Keeling, 1982; Silverman et al., 2010; Grimsley et al., 2011; Portfors, 2007). However, apart from representing the negative valence of social isolation, we speculated that USVs could also be studied during social reunion, and reflect the pups’ motivational drive to engage in social interaction (Matthews and Tye, 2019). To study maternally-directed behavior in mouse pups, we separated P15 mice from their dam and littermates for 3 hours and studied their USV emission and nipple attachment behavior upon reunion with an anesthetized dam. We observed that acutely-separated pups displayed a host of maternally-directed behaviors upon reunion, including robust USVs that were most prominent in close proximity to the dam (Fig. 1A-B). This finding is consistent with recent work in adult mice that observed isolation-induced increase in USVs during reunion (Zhao et al., 2021, 2023; Liu et al., 2023a), suggesting a context-dependent use of USVs also during mother-pup interaction. While MS did not affect pups’ engagement in nipple attachment behavior compared to unseparated pups (Fig. 1C-D, Extended Data Fig. 1A-D), we found that maternally-separated pups vocalized across varying attachment states-both outside and while engaging in nipple attachment (Fig. 1B), a phenomenon previously reported only in piglets but not in rodents (Jensen and Algers, 1984; Algers and Jensen, 1985). Using an unbiased approach for USV classification (Fig. 1J-K), our data revealed unique USV profiles that were linked to distinct states of attachment behavior. Specifically, MS pups emitted more USVs characterized by narrow bandwidth and high mean frequency (”Narrow-High”) outside of attachment events, compared to USVs with narrow bandwidth and low mean frequency (”Narrow-Low”) that were more common during attachment (Fig. 1L-N). Previous studies reported that USVs in rat pups are associated with different aspects of maternal care (Boulanger-Bertolus et al., 2017; Hofer, 1996). Recently, Opendak et al. showed that the dam’s presence buffered USVs in infant rats exposed to acute shock (Opendak et al., 2021). While our findings correspond with these maternal effects, they distinctly suggest a homeostatic framework underlying spontaneous vocal communication in the pre-weaning period, and emphasizes nipple attachment behavior as a potential regulator of USV emission during mother-pup interaction.

The social homeostasis theory has received much attention in recent years, pointing to OT as a key driver of isolation-induced social motivation (Matthews and Tye, 2019). While much of the literature has focused on the effects of chronic social isolation on the OT system (Veenema, 2012), few studies have delved into the functional responses of OT to acute social isolation at any age, much less in early life. We found that hypothalamic OT neurons increased their activity during MS, and returned to baseline upon reunion (Fig. 2A-B), consistent with previous work in P9 prairie voles showing elevated OT immunoreactivity after acute isolation (Kelly et al., 2018). We further showed that the density of activated OT neurons was higher in the caudal PVN, and that this activation was not coupled with elevated peripheral secretion of OT (Fig. 2D-E). These findings suggest a selective role of parvo-OT neurons in promoting MS-induced maternally directed behavior during reunion. This hypothesis is further supported by a recent study showing that the majority of activated OT neurons during acute social isolation in adult mice are enriched in *Cnr1* (Liu et al., 2023a), a molecular marker of parvo-OT neurons (Lewis et al., 2020). We excluded the possibility that these results were due to underdevelopment of OT neurons at this age, by demonstrating that magno-OT and parvo-OT subpopulations in the PVN are anatomically and functionally mature as early as the second week of life in mice (Extended Data Fig. 3; Luther and Tasker, 2000; Lewis et al., 2020). Parvo-OT neurons may contribute to maternally-directed behavior through local modulation of the larger magno-OT subpopultion (Tang et al., 2020b; Eliava et al., 2016) or through synaptic release of OT in specific targets like the nucleus accumbens (NAc) and periaqueductal gray (PAG), brain regions that receive synaptic inputs of parvo-OT from the PVN and regulate social reward and USV emission, respectively (Lewis et al., 2020; Iwasaki et al., 2023; Chen et al., 2021a). In addition, it has been recently found that separation calls and nipple attachment are independently regulated by agouti-related peptide (Agrp) neurons in the arcuate nucleus (Zimmer et al., 2019), proposing, together with our findings, a downstream role of parvocellular OT neurons (Atasoy et al., 2012) in controlling maternally-directed behavior.

In support of a causal role for OT in MS-induced behaviors, antagonism of OT receptors in separated pups reduced nipple attachment behavior and altered attachment-associated modulation of USVs upon reunion (Fig. 3, Extended Data Fig. 5). These results are consistent with recent work showing that OT and OTR knockout rat pups had impaired preference for home-cage bedding over unfamiliar bedding, but distinct in that knockout rats were not defective in USV emission during MS (Liu et al., 2023b). Although the different models (i.e., pharmacological vs. transgenic) and contexts (i.e., reunion vs. separation USVs) are difficult to compare, the inter-species variability in OTR developmental expression might contribute to the species- and context-specific behavioral repertoire and explain the different responses to OTR perturbations (Grinevich et al., 2014; Jurek and Neumann, 2018). Additionally, in our study silencing of OT neurons during MS did not affect the overall USV counts emitted during separation, but rather changed the dynamics of USV emission over the course of 3-hour inhibition (Fig. 5B-C), suggesting a complex and nuanced role of OT signaling in modulating separation-induced USVs in early life. This may help to clarify previous inconclusive findings regarding the role of OT in MS-induced USVs (reviewed in Hammock, 2015), underscoring the importance of extended observation periods to fully understand the subtleties of OT’s effects on early-life vocal communication.

To study the OT-mediated behavioral responses in pups during separation and reunion with higher spatial and temporal precision, we developed an optogenetic approach for cell type-specific silencing in freely-behaving pups. Recent advances in optogenetic technology have allowed minimally-invasive modulation of deep neuronal populations with high temporal and cellular specificity (Chen et al., 2021b; Chuong et al., 2014; Gong et al., 2020; Lin et al., 2013). These strategies are essential in the study of the developing brain, where intracranial implantation is technically difficult, and can adversely and unpredictably affect typical neurodevelopment (Xu et al., 2007; Polikov et al., 2005). However, most of these strategies were aimed for experiments that require neural excitation, and all of these studies still required the implantation of optical fibers on the dura or directly on the skull. To overcome this, we used eOPN3, a potent and highly light-sensitive inhibitory opsin (Mahn et al., 2021), to establish transcranial, deep and prolonged photoinhibition in unthetered, freely-behaving pups. Unlike other inhibitory optogenetic tools that act as ion channels or pumps (Mahn et al., 2016; Wiegert et al., 2017), eOPN3 is a light-activated inhibitory G-protein coupled receptor (GPCR), capable of prolonged signal transduction after activation (Koyanagi et al., 2013) and is thus suitable for manipulations that require vesicle release inhibition that recovers within minutes (Mahn et al., 2021). While chemogenetic silencing is also feasible in pups, the temporal precision of this approach is lower (recovery within the hours timescale; Wiegert et al., 2017) and it requires direct injection of the agonist, which can increase stress and interfere with behavioral readouts. We found that eOPN3 can effectively suppress OT neuron activity with low-irradiance red light (Fig. 4D-J), making it a suitable tool for transcranial inhibition of deep brain structures like the PVN. These properties of eOPN3 allowed us to design a minimally-invasive surgical procedure and a custom red LED array for home-cage manipulations *in vivo* (Extended Data Fig. 6). We showed that, with this configuration, sufficient red light reaches the ventral surface of the brain to silence the neural activity of hypothalamic OT neurons in freely-behaving pups (Fig. 4K-O), paving the way to optogenetic silencing in pups at this early age.

Using this approach, we found that silencing of OT neurons in pups during MS prevented the natural attenuation of USVs during separation, and selectively disrupted vocal behavior during reunion (Fig. 5). Notably, while USV emission during separation strongly predicted vocal behavior upon reunion, this relationship was abolished when OT neurons were inhibited. These results are consistent with our pharmacological results, where application of OTRA during MS had profound effects on USV emission during reunion. This demonstrates the importance of OT signaling during MS, not only influencing vocalizations during separation period but also shaping vocal behavior upon reunion. However, while optogenetic silencing of OT neurons during MS had no effect on non-vocal behaviors during reunion (i.e., nipple attachment), OTRA administration did affect these behaviors. This contrasting effect might arise from the slow pharmacokinetics of OTRA (half-life of 2 hours; Thompson et al., 1997), compared with the minutes-scale recovery kinetics of eOPN3. Although OT neuron activity in separated pups was decreased upon reunion, this might be attributed to the inherent limitations of using c-Fos as a proxy for neural activity (Barros et al., 2015; Morgan et al., 1987).

There is no consensus in the literature regarding OT’s sex-specific effects (Caldwell, 2018). We showed sex differences in MS-induced vocal communication during reunion, but not during separation (Fig. 5), consistent with the well-established sex differences in spontaneous and context-dependent vocalizations previously reported in rodents and humans (Ey et al., 2018; Liu et al., 2023a; Zhao et al., 2021, 2023; Oller et al., 2023). Unexpectedly, we further found that the effect of OT neurons on MS-induced USV emission during reunion was sexually-dimorphic, revealing a specific effect in female pups but not in males. Several studies have recently found sexual dimorphism in OT-mediated effects on behavior in juvenile rats (Dumais and Veenema, 2016; Maroun et al., 2020). However, to the best of our knowledge, our results constitute the first demonstration of sex-specific differences in OT-mediated effects in the pre-weaning period. These early findings imply that the neural mechanisms by which OT regulates context-dependent vocal behavior might evolve differently in females and males. This could be explained by sex differences, either in the developmental distribution of OTR and OT fibers (Caldwell, 2018), in the electrical properties of OT neurons (Murai et al., 1998), or in the types and properties of neurons expressing OTR (Li et al., 2016; Nakajima et al., 2014). Together, these results indicate that a combination of sex- and context-dependent mechanisms drive the responses of OT and OT-sensitive networks (Scott et al., 2015; Dumais and Veenema, 2016; Steinman et al., 2016; Winter and Jurek, 2019) to modulate maternally-directed behavior in the pre-weaning period.

Our results reveal a unique role for OT neurons in driving context-dependent maternally-directed behavior in pre-weaning mice. Using pharmacology and non-invasive optogenetic interventions, we demonstrate that OT modulates the vocal behavior of mice during both separation and reunion. Our findings emphasize the need to study the mechanisms through which the OT system modulates context-specific behaviors during early life, and to gain a more nuanced understanding of complex social interaction at this age. The new tools described here will encourage further research of the neural circuit mechanisms of behavior during the pre-weaning age, allowing new insights into the development of social behavior in early life.

## Materials and Methods

### Animals

Animals used for this study were preweaning male C57BL/6Jmice (Envigo) and the following transgenic lines: Oxttm1.^1(cre)Dolsn^/J (OT-IRES-Cre; Jackson Laboratories; 024234; Wu et al., 2012); Cg-Gt(ROSA)26Sor^tm9(CAG-tdTomato)Hze^/J (Ai9 reporter line; Jackson Laboratories; 007909; Madisen et al., 2010). OT-Cre mice were maintained by breeding homozygous males with heterozygous females, and pups for the experiments (OT-Cre^+/-^) were generated by breeding of homozygous males with wild-type C57BL/6J females. For experiments in Fig. 4, Fig. 5, Extended Data Fig. 3, Extended Data Fig. 7, both males and females were used. For *in vivo* behavioral experiments, pups were cross-fostered with age-matched littermates at P0-P3 and housed in litters of 5-8 pups with ICR foster dams (Envigo). Littermates from single cages underwent surgeries on the same day and were assigned to the experiment or control group such that cages always included mixed groups. Mice were kept in temperature- and humidity-controlled rooms on a 12-hour light–dark cycle with food and water ad libitum. All behavioral experiments were conducted during the dark phase. Experiments at the Charité-Universitätsmedizin Berlin were approved by the Berlin local authorities and the animal welfare committee of the Charité-Universitätsmedizin Berlin, Germany, and were done according to the guidelines stated in Directive 2010/63/EU. All procedures described in this paper were approved by the Weizmann Institute Animal Care and Use Committee.

### Mother-pup assay

The mother-pup (MP) assay was employed to study maternally-directed behavior in P15-P16 mouse pups. This behavioral paradigm allowed us to examine pup behavior toward an anesthetized dam after a 3-hour maternal separation (MS) period. The design of this assay was inspired by previously described behavioral assays (Zimmer et al., 2019; Opendak et al., 2021), with a few modifications. Notably, our assay included a 3-hour MS phase followed by a synchronous assessment of pups’ USVs and physical behavior toward the dam during a 10-minute reunion phase. All pups that underwent the MP assay were cross-fostered with an ICR lactating dam at the same postpartum age. Cross-fostering occurred immediately after neonatal injections at P0 (Fig. 5) or at P2-P3 for naive or pharmacologically-treated pups (Fig. 1, Fig. 3). Utilizing ICR foster dams, widely used as recipient dams, significantly facilitated pups’ survival, and also enhanced the contrast ratio of video recording during mother-pup interaction (i.e., dark pup on a white dam). During the MS phase, P15-P16 pups were transiently separated from their dam and littermates and individually placed in a small cage with fresh bedding for 3 hours. To maintain the pups’ body temperature, cages were transferred to a separate room and placed on heating pads set to 32*◦C*. Aged-matched, unseparated control pups were removed with their dam and littermates to a new cage with fresh bedding and placed back on the rack for 3 hours. After 3 hours, pups were monitored during a 10-minute behavioral test (i.e., reunion) in a new cage (501 cm^2^ floor area; Tecniplast; GM500), covered with soiled bedding from their home cage, with an anesthetized ICR dam (100 mg/kg Ketamine, 10 mg/kg Xylazine; i.p.) placed in the opposing corner of the pup’s starting point with her nipples facing up. We used anesthetized dams to eliminate any participation of the dam in the behavior, and focus our investigation on the pups’ behavior. All behavioral experiments were conducted during the dark phase by a trained experimenter who was blinded to the mouse identity.

### Ultrasonic vocalization (USV) recording and quantification

USVs emitted by the pups were recorded in a sound-attenuating cubicle (Med Associates) equipped with an ultrasound-sensitive microphone (CM16/CMPA; Avisoft-Bioacoustics) situated *≈* 30 cm above the cage surface. Data were acquired at a sampling rate of 250 kHz and 16-bit resolution using an UltraSoundGate 416H and an Avisoft-RECORDER software (Avisoft Bioacoustics). Gain was manually adjusted online to prevent signal saturation. USVs were detected using DeepSqueak version 2.6.2 (Coffey et al., 2019) using the ”All Short Calls” v1 network with the following parameters: high recall, ”Chunk length” = 5 sec, ”Overlap” = 0.06 sec. We excluded frequencies below 25kHz and above 120 kHz, and detected signals *>* 150 ms to reduce non-USV noise and artifacts. The performance of DeepSqueak was quantified using the overlap between ground-truth (GT) and predicted USVs (i.e., intersection-over-union). Bounding boxes for individual USVs were manually annotated (n = 802 USVs in 2 mice) by a trained observer and served as GT. Best-predicted fit: median = 0.905, IQR = [0.763 − 0.963]; best-GT fit: median = 0.884, IQR = [0.664 − 0.958].

We employed two classification methods for the detected USVs. First, we used VocalMat classifier (Fonseca et al., 2021), which uses a supervised learning model to classify the detected calls into a predefined set of vocal classes (adopted from Scattoni et al., 2008; Grimsley et al., 2011). We evaluated VocalMat performance by comparing the most likely class predicted by the classifier to the class assigned by a trained observer (n = 1950 in 6 mice; Extended Data Fig. 2B). We maximized the overall F1 score of the model by calculating the optimal prediction score (i.e. threshold; [0-1]) that optimizes F1 measure for each category. In addition, we expanded the analysis to consider one of the two most likely USV types assigned by the model, based on a classification approach of multi-class probability (Fonseca et al., 2021). This strategy improved the overall accuracy of the model (overall F1 score = 50.3%), mainly for the more prevalent categories. Based on this analysis, we conservatively included in all further analyses vocal classes that could be reliably predicted by VocalMat (i.e., accuracy *>* 50%; *chevron, down fm, flat, short, step up, up fm*). USVs that could not be classified using this method were labeled as *other*.

Additionally, we used the bandwidth and mean frequency of individual USVs to conduct an unsupervised classification of the detected USVs using k-means cluster analysis. Data were standardized to Z-scores and *kmeans* function was employed in MATLAB with 10 replicates to search for a local minimum. To determine the optimal number of clusters for this classification, we employed silhouette analysis using the *silhouette* function in MATLAB. This analysis was done separately for each experimental group. Mice with less than one USV per cluster were excluded from the analysis.

### Video recording, synchronization and quantification

Top-view videos of the reunion phase were recorded at 25 frames per second (1280 X 1024 resolution) using a GigE monochrome camera (DMK 33GP1300, Imaging Source) attached to a lens (H3Z4518CS-MPIR, Computar) under 850-nm IR illumination for 10 minutes. Videos were recorded using Streampix8 (Norpix, Canada), coupled to a custom Arduino-based system that allowed the acquisition and control of four independent setups simultaneously. An Arduino-controlled LED (1 W, 820 nm) was placed in each camera’s field of view and flashed at 0.05 Hz (100 ms pulse width) to facilitate the synchronization of video frames with the audio signal. A TTL-based signal, driven by the same pulse used for the video sync signal, was encoded in the least significant bit of the 16-bit audio sample and synchronized to the video stream via a custom MATLAB script. Automated tracking of recorded videos was performed using a custom MATLAB script, using frame-by-frame threshold-based segmentation and identification of the centroid. Manual analysis of behavior (i.e., nipple attachment events) was performed using a custom MATLAB script. All analyses were conducted by a trained observer blinded to the mouse identity.

### Analysis of USV emission and its relation to nipple attachment behavior

For the analysis of short- and long-range USVs (Fig. 1I, Extended Data Fig. 4J), we defined the “mother-zone” as a 7-cm radius from the dam’s center of mass, setting a threshold for short-range (i.e. within the mother-zone) and long-range (i.e. outside the mother-zone) USVs. We analyzed the USV rates of short- and long-range USVs before the first nipple attachment.

The ”USV-attachment bias” index (Fig. 1L, Fig. 3D,I,J) was calculated per mouse for total USV counts or specifically for the three USV clusters (i.e., k-means cluster analysis) as:

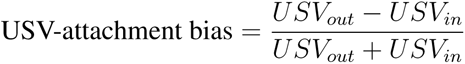

where *USV_out_* is the USV rate outside nipple attachment and *USV_in_* is the USV rate within nipple attachment. Mice that did not display nipple attachment for the entire trial were excluded from the current and subsequent analysis.

For the analysis of cluster-specific USV emission in relation to the first nipple attachment (Fig. 1M, Extended Data Fig. 5C,G), we calculated the Z-score distribution of USV rate per mouse for each cluster over 60-second time bins. These time bins were aligned to the time of first nipple attachment and averaged across mice. We considered only time bins that contained data from all mice for the statistical analysis examining the effect of cluster and time from first nipple attachment on cluster-specific USV emission.

In addition, we used the ”USV-1st attach bias” index to quantify cluster-specific tendency to vocalize in relation to first nipple attachment (Fig. 1N, Extended Data Fig. 5D,H). This index was calculated as the difference between the mean Z-scores over 60-sec time bins before and after first nipple attachment per mouse, and then averaged these values across mice for each cluster. We included all time bins in this analysis per mouse.

To study the temporal dynamics of USV emission in relation to the onset of nipple attachment (Fig. 3E,G,F,H), we pooled together attachment events for each experimental group and analyzed the USV rates aligned with the onset of events. The USV data were discretized into 100-ms bins, and smoothed using a moving average (window size = 30 bins). Peri-event time histograms (PETHs) were averaged across attachment events, both for total USV counts and for the three specific USV clusters, spanning from -20 to +20 seconds from nipple attachment, and binned at 100 ms intervals. The results were also superimposed with the mother-pup center-of-mass distances, calculated in 100-ms bins and smoothed using a moving average (window size = 30 bins).

### Pharmacological study *in vivo*

Oxytocin receptor antagonist (OTRA; L-368,899; Tocris 2641) was dissolved in saline 95% and DMSO 5%. We applied 10 mg/kg OTRA intraperitoneally 30 minutes before maternal separation in a volume of 10 ml/kg mouse weight. Control animals were injected with vehicle solution containing saline 95% and DMSO 5%. Mice were assigned randomly to OTRA or control groups and litters contained mixed groups.

### Primary dissociated hippocampal neuron culture and gene delivery

Autaptic primary hippocampal neuronal cultures on glial cell micro-islands were prepared from P0 mice (C57BL/6NHsd; Envigo) of either sex as previously described (Rost et al., 2010). 300 µm diameter spots of growth-permissive substrate consisting of 0.7 mg ml-1 collagen and 0.1 mg ml-1 poly-D-lysine were applied with a custom-made stamp on agarose-coated coverslips. First, astrocytes were seeded and allowed to proliferate in Dulbecco’s modified eagle medium (DMEM) supplemented with 10% fetal calf serum and 0.2% penicillin/streptomycin (Invitrogen) for one week to form glia micro-islands. After changing the medium to Neurobasal-A supplemented with 2% B27 and 0.2% penicillin/streptomycin, hippocampal neurons prepared from P0 mice were added at a density of 370 cells/cm^2^. rAAV2/1 particles expressing eOPN3-mScarlet under the CaMKII*α* promoter were produced in accordance with the protocol described in (Mahn et al., 2021). Neurons were transduced with rAAV2/1-CaMKIIa-eOPN3-mScarlet (1.5 × 10^8^ VG per well) at DIV 1 and were recorded between DIV 14 and DIV 21.

### In vitro electrophysiology

EPSCs from autaptic primary neurons were recorded under visual guidance using an Olympus IX51 inverted microscope with an Olympus UPlanSApo 20x/0.75 UIS2 objective under infrared widefield illumination. A CoolLED P4000 served as a light source to identify expressing cells and for light activation of eOPN3. To avoid pre-activation of eOPN3 by fluorescence excitation light, electrophysiological recordings were performed first, and cells were only then investigated for expression. Acquired data was excluded in case cells were not expressing. The activation light was filtered with a narrow bandpass filter (center wavelength 636 ±10 nm, #65-106, Edmund Optics) and additionally attenuated with a 6.0 neutral density filter. Light intensities were measured with a calibrated S130VC power sensor (Thorlabs). Autaptic neurons were constantly perfused with extracellular solution (in mM): 140 NaCl, 2.4 KCl, 10 HEPES, 10 D-glucose, 2 CaCl_2_, and 4 MgCl_2_ (pH was adjusted to 7.3 with NaOH, 300 mOsm). Cells were patched with microelectrodes pulled from quartz glass capillaries (3-4 MΩ), filled with (in mM): 136 KCl, 17.8 HEPES, 1 EGTA, 0.6 MgCl_2_, 4 MgATP, 0.3 Na_2_GTP, 12 Na_2_ phosphocreatine, 50 U/ml phosphocreatine kinase (300 mOsm); pH adjusted to 7.3 with KOH. A Multiclamp 700B (Molecular Devices) amplifier and NI USB-6343 digitizer (National Instruments) were used to control and acquire electrophysiological recordings and the application of light stimulation via WinWCP 5.7 software (https://github.com/johndempster/WinWCPXE). Data was acquired at 10 kHz and filtered at 3 kHz. Cells were kept at -70 mV, and series resistance and capacitance were compensated by 70%. EPSCs were elicited by a 1 ms depolarization to 0 mV (50 ms interstimulus interval, every 5 s), resulting in an unclamped axonal action potential causing neurotransmitter release. Experiments were performed at room temperature. Data were analyzed using Clampfit 10.7 (Molecular Devices). Cells were excluded from analysis if the first EPSC amplitude was below 100 pA, if the access resistance was above 20 MΩ, or if the holding current exceeded 200 pA. All data points represent measurements from biological replicates.

### Acute slice electrophysiology

Recordings were performed from acute coronal slices prepared from pups between P14 and P21 (see below). For recording OT neuron activity after maternal separation and electrophysiological identification of magnocellular and parvocellular OT neurons (Extended Data Fig. 2A-F), we used an OT-IRES-Cre driver line crossed to a Cre-dependent TdTomato reporter line (OT-IRES-Cre x Ai9). For eOPN3 activation measurements (Fig. 4D-K), we used OT-IRES-Cre pups injected at P0 into the PVN with a Cre-dependent rAAV2/1 vector expressing eOPN3 (pAAV-hSyn1-SIO-eOPN3-mScarlet-WPRE; Addgene #125713).

### Acute brain slice preparation

Pups were injected with pentobarbital (100 mg/kg, i.p.). After decapitation, 300 µm-thick coronal slices containing the PVN were prepared in carbogenated (95% O_2_, 5% CO_2_) ice-cold slicing solution containing (in mM): 2.5 KCl, 11 glucose, 234 sucrose, 26 NaHCO_3_, 1.25 NaH_2_PO_4_, 10 MgSO_4_, 2 CaCl_2_; pH 7.4, 340 mOsm. Slices were cut using a vibratome (Leica VT 1200S) and allowed to recover for 20 min at 33°C in carbogenated high osmolarity artificial cerebrospinal fluid (aCSF, high-Osm) containing (in mM): 3.2 KCl, 11.8 glucose, 132 NaCl, 27.9 NaHCO_3_, 1.34 NaH_2_PO_4_, 1.07 MgCl_2_, 2.14 CaCl_2_; pH 7.4, 320 mOsm. Subsequently, slices were incubated for 40 min at 33°C in carbogenated aCSF containing (in mM): 3 KCl, 11 glucose, 123 NaCl, 26 NaHCO_3_, 1.25 NaH_2_PO_4_, 1 MgCl_2_, 2 CaCl_2_; pH 7.4, 300 mOsm. Finally, slices were kept at room temperature (23°C –25°C) in the same solution until use.

### Whole-cell patch-clamp recording

OT neurons in the PVN were patched under visual guidance using infrared differential interference contrast (DIC) microscopy (BX51W1, Olympus) and an Andor Neo sCMOS camera (Oxford Instruments). Borosilicate glass pipettes (BF100-58-10, Sutter Instrument, Novato, CA, USA) with resistances 4–6 MΩ were pulled using a laser micropipette puller (P-2000, Sutter Instrument) and filled with intracellular solution (in mM): 135 potassium-gluconate, 4 KCl, 2 NaCl, 10 HEPES, 4 EGTA, 4 Mg-ATP, 0.3 Na_2_-GTP, 10 phosphocreatine-Na_2_, 280 mOsm, pH adjusted to 7.3 with KOH. Somatic whole-cell voltage-clamp recordings (¿1 GΩ seal resistance, -70 mV holding potential) were performed using a Multiclamp 700B amplifier (Molecular Devices). Data were acquired using pCLAMP 10.7 on a personal computer connected to the amplifier via a Digidata-1440 interface (sampling rate: 20 kHz; low-pass filter: 4 kHz) and analyzed with Clampfit 10.7 (all Molecular Devices). Data obtained with a series resistance ¿ 20 MΩ were discarded. All experiments were conducted at room temperature. In the recording chamber, slices were superfused with carbogenated aCSF (4–5 mL/min flow rate). Data were analyzed using Clampfit 10.7 (Molecular Devices).

### OT neuron activity after maternal separation

We tested the effect of a 3-hour MS on PVN-OT neurons intrinsic membrane properties (n = 19) as well as on their excitatory and inhibitory synaptic inputs (sEPSCs: n = 18; sIPSCs: n = 17 in 5 mice). We used aged-matched undisturbed pups as controls (intrinsic properties: n = 14; sEPSCs: n = 14; sIPSCs: n = 13 in 4 mice). PVN-OT neurons were identified by the endogenous expression of tdTomato (OT-IRES-Cre x Ai9). As for intrinsic membrane properties, the membrane resistance (Rm), time constant (*τ* m), rheobase and cell excitability were measured. For this, OT neurons were recorded in current-clamp mode at a resting potential of -65 mV. A slow current ramp was applied to the cell to calculate the rheobase, while for Rm, *τ* m and cell excitability, a succession of square pulses was applied (from -20 pA to +40 pA, 5 pA steps). Rm and *τ* m were measured during hyperpolarizing pulses, while the number of AP was plotted for each depolarizing step for cell excitability. Afterwards, spontaneous excitatory (sEPSCs) and inhibitory (sIPSCs) postsynaptic currents were recorded in voltage clamp mode (holding -70 mV for sEPSCs and 0 mV for sIPSCs), and the amplitude and frequency were measured.

### OT-eOPN3 activation measurements

PVN-OT neurons were identified by the viral expression of mScarlet (AAV-hSyn-SIO-eOPN3-mScarlet). Cells were recorded in current clamp mode at their resting membrane potential if spontaneously active. In cells that did not spontaneously spike, a small constant depolarizing current was applied to reach a stable low firing rate. Light was delivered using a Lumencor SpectraX light engine, using band-pass filters at 572/35 and 632/22 nm (peak wavelength/bandwidth). The effect of eOPN3 activation on neuronal firing by a long (60 sec) pulse of low intensity red light (632 nm at 47 µW/mm^2^) was compared to the previously described illumination protocol (Mahn et al., 2021) with a brief (500 ms) pulse of green light (572 nm at 365 µW/mm^2^). For the comparison of post-illumination firing rate in OT-eOPN3 neurons, the values presented are normalized to mean firing rate within a 30-s baseline period and averaged across cells (Fig. 4F,I).

### Classification of magno-OT and parvo-OT neurons

For the classification of magno-OT and parvo-OT neurons, we used both the morphological features of the cells and the electrophysiological protocol previously described (Lewis et al., 2020; Eliava et al., 2016). Briefly, each cell was hyperpolarized to -100 mV and then depolarized using current steps of increasing amplitude. To discriminate between magno-OT and parvo-OT neurons, we measured the full width at half maximum of the first action potential (”AP duration”) and the latency to the first action potential.

### Stereotaxic injection of AAV vectors

Neonatal stereotaxic injections were conducted as described previously (Kim et al., 2014), with a few modifications. Due to the lack of a standard procedure for neonatal injections into the PVN, we first performed a pilot study to determine the optimal coordinates for PVN injections in newborn mice, as well as the viral serotype and volume of virus required for appropriate viral expression in OT neurons (data not shown). These parameters were used in the following procedure. Briefly, newborn pups were injected after birth (P0), after they had started nursing (6-12 hours after parturition). Pups were anesthetized via hypothermia and placed in a stereotaxic apparatus (David Kopf Instruments) using a custom neonatal stage with soft ear bars and a reservoir for ice to maintain anesthesia throughout the procedure. To avoid direct contact with ice, pups were covered with a piece of a nitrile glove. The sagittal and transverse sinuses, clearly visible through the skin at this age, were used as landmarks for our injections, and we defined their intersection as lambda. The anesthetized pup’s head was disinfected with 70% ethanol. A Nanofil syringe (World Precision Instruments) with a 34G beveled needle was filled with AAV suspension. The needle was inserted through the scalp and skull into the PVN, bevel facing anterior, and left in place for 1 minute, followed by slowly injecting 300 nL of the virus (100 nL/min). After injection, the needle was left in place for an additional 2 minutes and then slowly withdrawn. PVN coordinates in mm relative to lambda: AP: 2.3, ML: *±* 0.35, DV: -2.9. The DV coordinate was zeroed under the skin. The duration of the entire procedure did not exceed 30 minutes. After completing injections in both hemispheres, pups were placed on a heating pad and monitored until their skin color and movement returned to normal. Then, pups were marked with a paw tattoo for identification (Ketchum Manufacturing Inc.). After recovery, pups were cross-fostered with experienced ICR dams, widely used as recipient dams, and closely monitored for their well-being.

Adeno-associated viruses (AAV) used in this study were purchased from Addgene. The titters of AAV vectors used for intracranial injections are *after* dilution in sterile PBS x1 and presented in genome copies per milliliter (gc/ml). AAV vectors in this study included: AAV1-hSyn1-DIO-EGFP (7.3 × 10^12^ gc/ml; plasmid #50457), AAV1-hSyn1-SIO-eOPN3-mScarlet-WPRE (7.3 × 10^12^ gc/ml; plasmid #125713). For all experiments, pups were tested at least 14 days after viral injections.

### Intact-skull implantation and transcranial illumination

For transcranial activation of eOPN3, we designed a minimally-invasive surgical procedure that allows optical access to the pups’ brain using a 5 mm diameter glass implanted onto the intact skull. P12-P13 pups were removed from the home-cage and placed on a heating pad while awaiting the surgery. Pups were anesthetized with a combination of anesthetic drugs (Medetomidine 0.5 mg/kg; Midazolam 5 mg/kg; Fentanyl 0.05 mg/kg; i.p.), a well-established anesthetic technique in neonatal mice (> *P* 2; Tang et al., 2020a). Once the pups were fully sedated, ophthalmic ointment was applied, the hair on the scalp was removed, and pups were immobilized in a Kopf stereotaxic apparatus using modified soft ear bars. The scalp was disinfected with ethanol 70%. A midline incision (8 mm long) was performed in the skin to allow an access to the skull, starting from the anterior edge of the ears (above lambda) to the posterior part of the eyes. Circular excisions were made over the left and right hemispheres of the skull, and the exposed skull was cleaned with sterile saline (NaCl 0.9%) and dried off with a cotton-tipped applicator. Then, a 5 mm glass implant was glued to the exposed skull with cyanoacrylate (Krazy-Glue) and centered over the skull midline with posterior edges around lambda. C&B Metabond (clear; Parkell) was applied to the outside perimeter of the window, covering any exposed bone to ensure a complete seal with the surrounding skin and muscles. The implant is composed of two layers of round coverglass (5 mm, no. 1 thickness; Hecht Assistent) glued together with Norland Optical Adehesive No. 81. (NOA 81; Edmund Optics). After the glues are completely dry, the pups are administered with anesthetic antagonists (atipamezole 2.5 mg/kg; flumazenil 0.5 mg/kg; naloxone 1.2 mg/kg; s.c.) and postoperative analgesia (Carprofen 5 mg/kg; s.c.) and removed back to the heating pad. The entire procedure did not exceed 20 minutes. Once the pups fully recovered, they were returned to their dam and closely monitored until the experiment.

To measure eOPN3-induced changes in PVN-OT neuron activity *in vivo*, we delivered red light through the intact-skull implant of P15-P16 pups using two different light sources placed above the implant. For experiments that involve measurements in anesthetized pups (0.5%−1% isoflurane; Extended Data Fig. 7B-D), an optical fiber (500 µm diameter, NA 0.73) was placed above the implant, and pups were illuminated with red LEDs (630 nm; 150 *µ*W/mm^2^ at the fiber tip; Prizmatix). For experiments that involve measurements in freely-behaving pups (Fig. 4N-P), a custom 64-Watt red LED array (peak: 655 nm; 20 cm X 10 cm; Zeal-Tech Industries, India; Extended Data Fig. 6B) was installed at a distance of 20 cm above a clear plexiglas cage (10 cm X 8 cm X 14 cm) with a single mouse. In both experiments we used a continuous illumination protocol for 80 minutes. Five minutes after red light activation, pups were administered with hyper-osmotic solution (salt-loading; see below), and 90 minutes later, pups were euthanized for histological examination. Importantly, with these conditions, we did not observe any evidence of overheating and damage to the brain tissue.

We used the 64-Watt LED array to evaluate the amount of light that could travel through the tissue to reach the PVN. P15-P16 implanted pups were euthanized and decapitated. We removed the upper palate and carefully dissected out the pituitary gland to expose the ventral hypothalamus, where a light sensor (3 mm diameter; Thorlabs) was placed for further measurements. The red LED array was installed 20 cm above the testing surface and we calculated light transmission through the following tissue samples: intact-skull implant with brain and skull, brain and skull (i.e., skin removed), extracted brain only, skin with fur and skull only. To prevent false detection from ambient light and reflections, tissue samples were covered with aluminum foil, leaving a 5 mm window around the implant. The transmission percentage (T%) of red light was calculated using the formula 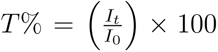, where *I_t_* is the intensity of light after passing through the tissue and *I*_0_ is the baseline light intensity without any tissue between the sensor and light source. Notably, the light power measured at ventral surface of the brain corresponded with an irradiance of 75 *µ*W/mm^2^ delivered by a fiber-coupled red LED (630 nm; Prizmatix) placed right above the implant. This light power is well tolerated by brain tissue and is safe for in vivo experiments (Cardin et al., 2010).

### Salt loading

We used acute salt loading as a method to induce robust c-Fos expression in hypothalamic OT neurons, a well-established procedure as previously described (Hasan et al., 2019; Hoffman et al., 1993). To validate the effectiveness of this approach in pups, P15 pups were injected with hyper-osmotic solution (NaCl 9%; 20 ml/kg mouse weight, i.p.) and compared to pups injected with normal saline (NaCl 0.9%). Pups were sacrificed 90 minutes after salt loading, and we assessed the expression patterns, as depicted in Fig. 7A. Subsequently, we applied this approach in the experiments aimed at establishing trasncranial photoinhibition using eOPN3 (Fig. 4N-P, Extended Data Fig. 7B-G). This procedure served as an important component of our experimental design, allowing us to reliably induce c-Fos expression in a high fraction of the OT population for subsequent investigations.

### Transcranial photoinhibition of OT neurons in behaving pups

To investigate the impact of eOPN3-induced inhibition of OT neurons during MS on pups’ behavior, AAV vectors encoding a Cre-dependent eOPN3-mScarlet transgene (rAAV2/1-hSyn1-SIO-eOPN3-mScarlet-WPRE; 7.3 × 10^12^ gc/ml; n = 19) or EGFP (rAAV2/1-hSyn1-DIO-EGFP; 7.3 × 10^12^ gc/ml; n = 26) were bilaterally injected into the PVN of OT-IRES-Cre P0 pups. At P12-P13 pups were implanted with an intact skull implant, and carefully monitored until the testing day. To minimize the incidence of aggressive behavior and infanticide by the dam towards the operated pups, dams were desensitized to the surgical substances starting 24-48 hours before the procedure (Cruz-Martin and Portera-Cailliau, 2014). At P15-P16 pups underwent MS in a clear plexiglass cage (10 cm X 8 cm X 14 cm) and were subjected to 3-hour illumination using an Arduino-triggered red LED array (655 nm; 1 sec on, 1 sec off protocol) positioned above the cage. During MS we installed a CM16/CMPA microphone (Avisoft) 20 cm above the cage surface, and sampled 5-min USV recordings at 20-min intervals, for a total of 8 recording sessions. Summary of USV count during MS, calculated as the median number of USVs during the first and second halves of MS per mouse and averaged across mice. Four pups were not recorded during MS and excluded from the current analysis (n_(OT-EGFP)_ = 24; n_(OT-eOPN3)_ = 17). Then, following a brief recovery period (5 minutes; red light off), pups were removed to a new cage with an anesthetized dam to assess behavior upon reunion, as detailed in the ”Mother-pup assay”.

To assess the impact of various features in the optogenetic experiment on pups’ behavior during the reunion phase, we fit a general linear model (GLM) with the following features:

- Mean fluorescence intensity (FI) of PVN sections for eOPN3- and EGFP-expressing pups. Values were averaged per mouse and standardized (Z-scores) per experimental group.
- MS USV attenuation index (U), computed using the following formula:

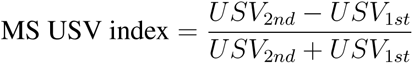

where *USV*_1*st*_ and *USV*_2*nd*_ are the median USV counts in MS first- and second-half recordings, respectively.
- Experimental group (g).
- Sex (s).

The predicting features showed low multicollinearity with a variance inflation factor *<* 1.4 for all variables.

For the predicted variables of the model, we categorized pup behavior upon reunion into USV-related data (”vocal behavior”): (1) USV rate until first attachment; (2) total USV count; (3) USV rate outside attachment events; and non-USV-related data (”non-vocal behavior”): (1) total time attached; (2) mean attachment length; (3) time to first attachment; (4) mean dam-pup distance; (5) entries to the dam’s zone. One pup was excluded from the PCA due to incomplete anesthesia of the dam and missing data during the reunion phase (n_(OT-EGFP)_ = 25; n_(OT-eOPN3)_ = 19). We then standardized these variables and projected the Z-scored values onto the first two principal components (PCs) using principal component analysis (PCA), that were considered two independent responses of the GLM.

Finally, we used the features described above to predict PC1and PC2 of pup behavior upon reunion. We removed pups where not all the predicting features could be measured (removing 7 pups where the mouse genotype was OT-Cre*^−/−^* and/or the mouse was not recorded during MS), as well as pups that were not recorded during reunion (removing 1 pup). This left n = 37 pups used for the model (EGFP: n_(F)_ = 10, n_(M)_ = 10; eOPN3: n_(F)_ = 8, n_(M)_ = 9). We used the *fitglm* function in MATLAB that received the following formula:

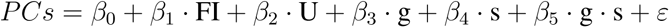

Where *PCs* represents PC1 or PC2, *β*_0_ is the intercept, *β*_1_ through *β*_5_ are the coefficients to be estimated for each predictor variable, and *ε* represents the error term. For post-hoc Bonferroni-corrected comparisons of estimated marginal means we used the *emmeans* function in R to compare the experimental group and sex.

### Blood sampling and quantification of plasma OT

Blood samples from the facial vein of P15 pups were collected in EDTA-coated tubes (Mini-Collect, De-Grut) 1 hour after MS or from unseparated control pups. Blood samples were centrifuged at 1300 *g* for 10 min at 4 *^◦^*C, and 100 µl plasma was collected for each mouse and stored at *−*80 *^◦^*C until radioimmunoassay was performed (RIAgnosis, Munich, Germany; Neumann et al., 2013). Briefly, samples were preincubated for 60 min with 50 µl rabbit anti-OT antibody, and then 10 µl 125I-labeled tracer (Perkin Elmer, USA) was added to each aliquot. After an incubation period of 3 days at 4 *^◦^*C, unbound radioactivity was precipitated by activated charcoal (Sigma Aldrich, USA). Under these conditions, an average of 50% of total counts are bound with ¡5% non-specific binding. The detection limit is in the 0.1 pg/sample range and the intra- and inter-assay variability is < 10%.

### Immunohistochemistry, imaging, and quantification

Mice were deeply anesthetized using pentobarbital (130 mg/kg; i.p.) and then transcardially perfused with ice-cold PBS (pH 7.4, 10 ml) followed by 4% paraformaldehyde (PFA, 10 ml) solution. Heads were removed and post-fixed overnight at 4 *^◦^*C in 4% PFA. Then, brains were extracted and transferred to 30% sucrose solution for at least 24 hours. Coronal sections (30 µm) were acquired using a microtome (Leica Microsystems) and preserved in a cryoprotectant solution (25% glycerol, 30% ethylene glycol in PBS, pH 6.7). Sections were washed 3 *×* 10 min in PBS, and incubated for 1 hour at room temperature (RT) in blocking solution: 20% normal horse serum (NHS), 0.5% triton-X100 in PBS. Primary antibodies were diluted in 2% NHS, 0.5% triton-X100 in PBS and incubated overnight at RT. The following antibodies were used: mouse anti-NP1 (OT staining; 1:2000; Merck; MABN844), rabbit anti-cfos (1:500; Santa-Cruz; sc-52), rabbit anti-cfos (1:2500; Cell Signaling; #2250). The following day, the sections were washed 3 *×* 10 in PBS, then incubated for 2 h at RT with the following fluorophore-conjugated secondary antibodies: goat anti-rabbit Alexa 488 (1:300; Abcam; ab150113), donkey anti-rabbit Cy5 (1:300; Jackson ImmunoResearch 711-175-152), goat anti-mouse Alexa 488 (1:300; ThermoFisher; A11029), donkey anti-mouse Cy5 (1:300; Jackson ImmunoResearch; 715-175-151), prepared in 2% NHS and PBS. Sections were washed 3 *×* 10 in PBS and stained with a nucleic acid dye (4,6-diamidino-2-phenylindole (DAPI), 1:10000). Finally, sections were mounted on gelatin-coated slides, dehydrated, and embedded in mounting medium (Aqua-Poly/Mount; Polysciences). Fluorescent images were acquired using a VS120 slide-scanning microscope (Olympus) at 10x or 20x magnification with DAPI, FITC and/or Cy3 and/or Cy5 channels.

The resulting images were then analyzed using ImageJ (2.14) to measure fluorescence within a specific region of interest (ROI). For unbiased ROI selection, in all experiments ROIs were identified using the DAPI channel and outlined based on anatomical landmarks in the Mouse Brain atlas (Paxinos and Franklin). The following analyses were done separately for each section, and individual values were averaged across sections of the target region per mouse, unless mentioned otherwise.

For images requiring c-Fos quantification (acquired with a 20x objective; 0.322^2^ microns/pixel), we employed the pretrained StarDist 2D model for automatic segmentation of c-Fos+ nuclei (probability threshold = 0.5, nms = 0.3; Schmidt et al., 2018) and counted particles using the Analyzed Particles algorithm (size: 250-1500 pixels; circularity 0.7-1) in imageJ. c-Fos counts are normalized to the ROI’s area (counts/mm^2^).

To assess colocalization of c-Fos immunoreactivity with OT+/eOPN3+/EGFP+ (”Y”) cells, we used StarDist segmentation previously applied for c-Fos analysis, and manually calculated the percentage of c-Fos+ Y+ cells out of the total number of Y+ cells within the ROI in both hemispheres. In addition, we divided the PVN into rostral (*−*0.58: *−*0.82 mm relative to bregma) and caudal (*−*0.94: *−*1.22 mm) sections and calculated the mean colocalization of c-Fos and OT immunoreactivity per PVN subarea. For images requiring evaluation of eOPN3 or EGFP viral expression within the PVN, a fixed rectangle ROI was defined outlining the PVN bilaterally (excluding the 3rd ventricle). Mean fluorescence values were measured for Cy3 and FITC channels,respectively, to represent the expression level of cells and projections within the ROI. One Re pup was excluded from the PVN analysis for technical reasons (related to Fig. 2). In all experiments, imaging acquisition parameters and the ensuing analysis pipeline were kept constant across mice, and experimenters were blinded to mouse experimental group during all steps that required manual annotations.

### Identification of magnocellular and parvocellular OT neurons in P15 pups

As described previously in adult mice, P15 pups were transcardially perfused 8 days after two intraperitoneal injections of FluoroGold (FG) over the course of 24 hours (1% in normal saline; 20 mg/kg; Santa Cruz; Sc-358883). Coronal sections of the PVN were stained with a primary antibody against FG (rabbit anti-FG; 1:100; Fluorochrome), and FG immunoreactivity was visualized by an Alexa 488-conjugated secondary antibody (goat anti-rabbit; 1:800; Abcam; ab150113). Expression patterns of FG and OT in the rostral-caudal axis of the PVN (−0.70:-1.22 mm relative to bregma) are shown in Extended Data Fig. 3G.

### Statistical analyses

Details of specific statistical designs and appropriate tests are described for each analysis in the appropriate figure legend and throughout the text. Unless otherwise stated, data are summarized as the mean *±* s.e.m., and single data points are marked on the appropriate figures, where n denotes the number of mice for behavioral trials and histological examinations, the number of USVs for cluster analysis and validation analysis of detection and classification tools, the number of nipple attachment events for PETH, and the number of cells for electrophysiological recordings. No statistical methods were used to predetermine sample sizes, but our sample sizes were chosen based on standards in the field. If applicable, data points excluded for any reason are detailed in the corresponding figure legend, and in the appropriate section in the Methods. Apart from the behavioral experiment in Fig. 1, experimenters were blinded to the mouse experimental group, viral vector, drug and sex during experimental sessions and during initial analyses of behavioral and histological data. To control for nested effects, we used linear mixed-effect models (LME) for statistical analysis and included mouse identity as a random effect in our model (Extended Data Fig. 1, Extended Data Fig. 4). All statistical tests presented in this manuscript are two-tailed. Significance was set at an alpha value of 0.05, and Bonferroni corrections were used when appropriate to correct for post hoc and multiple comparisons. Specific P values are detailed for each analysis in the corresponding figure legend, and in Supplementary Table 2, and significant comparisons are marked on the relevant figure panels. Levene’s test was used to assess the equality of variances, and statistical parameters were adjusted accordingly when needed. When applicable, the data distribution was assumed to be normal, but this was not formally tested. All analyses and subsequent statistical tests were performed using Matlab v.2018b (Mathworks), SPSS v.21 (IBM), R Statistical Software (R Project for Statistical Computing) and imageJ (2.14).

## Supplementary Information

**Extended Data Figure 1:**
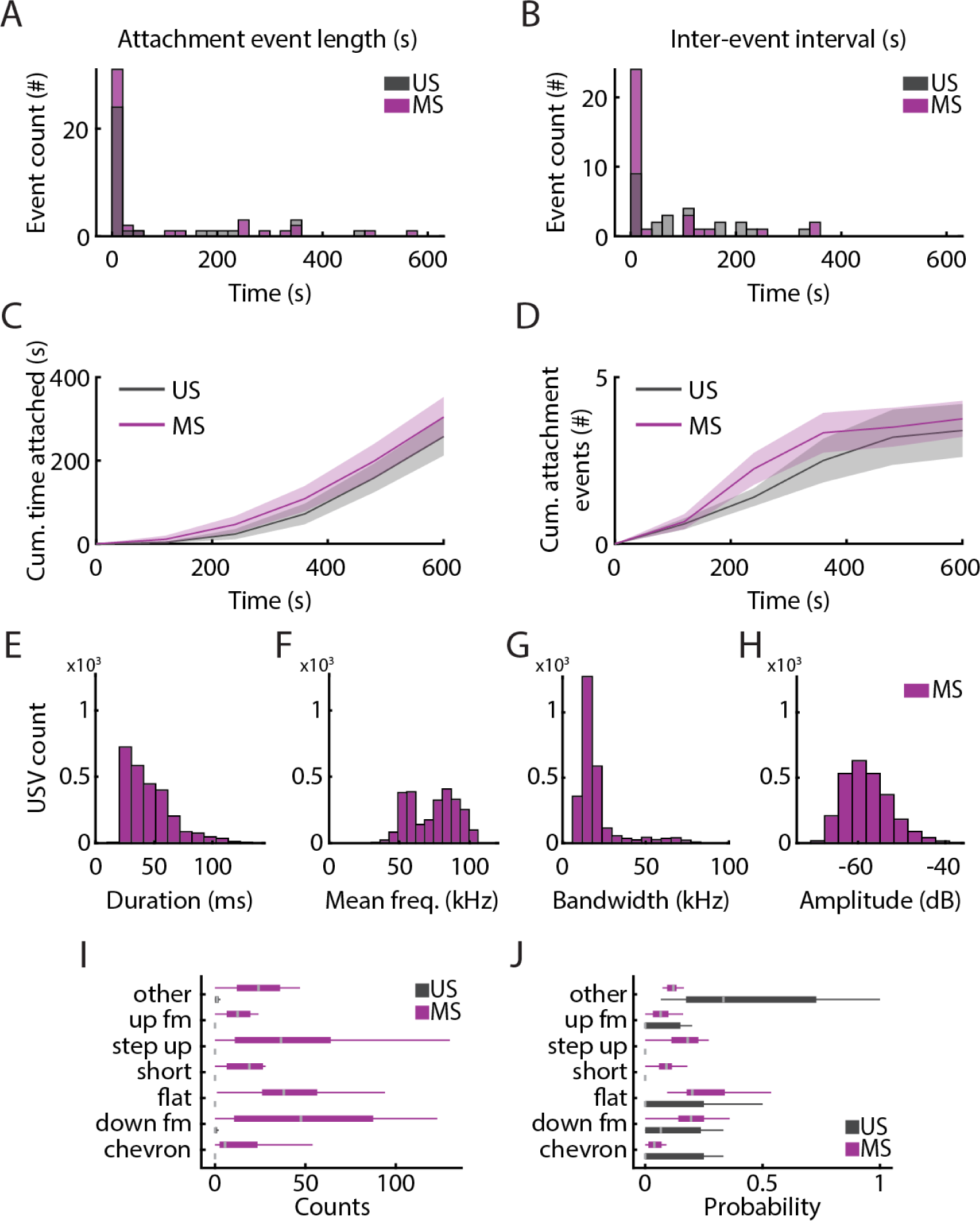
Maternal separation increases USV emission, but does not affect nipple attachment behavior in P15 pups. **Extended Data Figure 1: (A–B)** Distribution of (A) attachment event lengths and (B) interevent interval (IEI) over 20-sec bins for MS (n_(MS)_= 12; magenta) and US (n_(US)_ = 10; gray) pups in the MP assay. Fitting a linear mixed effect (LME) model for quantifying the differences in the distribution of events for MS and US pups. (A) *t*_(77)_ = 0.174, *P* = 0.863; (B) *t*_(55)_ = *−*1.523, *P* = 0.132. **(C)** Cumulative time of nipple attachment over 120-sec bins for MS and US pups. Mixed-design repeated measures ANOVA *F_time_*_(1.239,24.786)_ = 59.42, *P* = 1.1 *×* 10*^−^*^8^; *F_groups_*_(1,20)_ = 0.703, *P* = 0.412; *F_time×groups_*_(1.239,24.786)_ = 0.428, *P* = 0.561. **(D)** Cumulative counts of attachment events over 120-sec bins for MS and US pups. Mixed-design repeated measures ANOVA *F_time_*_(1.498,29.963)_ = 33.849, *P* = 1.64 *×* 10*^−^*^7^; *F_groups_*_(1,20)_ = 0.551, *P* = 0.466; *F_time×groups_*_(1.498,29.963)_ = 0.517, *P* = 0.55. Data in C,D are presented as mean *±* s.e.m. (shaded areas) **(E–H)** Properties of individual USVs emitted by MS pups, characterized by (E) duration, (F) mean frequency, (G) bandwidth and (H) amplitude. **(I-J)** Classification of USVs based on spectro-temporal properties, presented as (I) total counts and (J) probabilities of each vocal class averaged across MS and US pups. Box plot marks interquartile range (IQR; box edges) and median (gray line), whiskers mark 1.5*±* IQR.

**Extended Data Figure 2:**
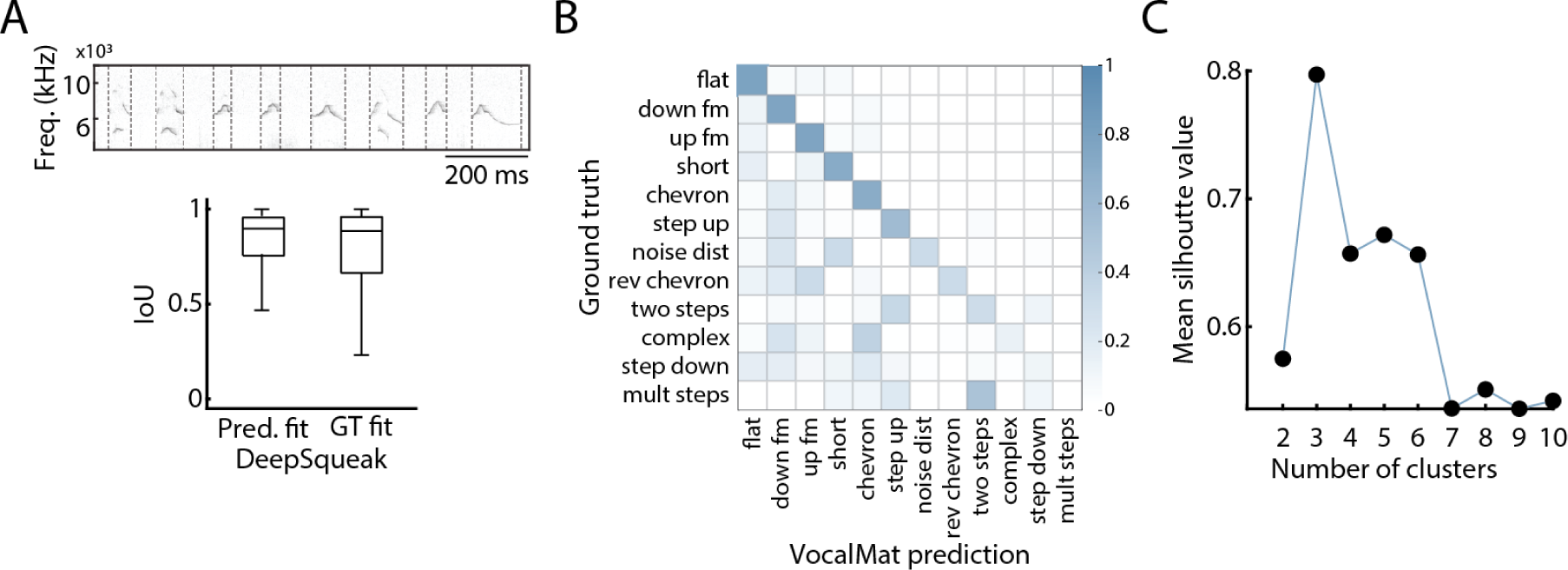
Detection and classification of USVs. **(A)** Top: Representative spectrogram of USV emission of MS pups upon reunion with the dam, showing bounding boxes for individual USVs (dashed line). Bottom: Validation of DeepSqueak performance. **(B)** Evaluation of VocalMat accuracy in classifying reunion USVs. **(C)** The optimal number of clusters used in the k-means cluster analysis was determined using the mean Silhouette value.

**Extended Data Figure 3:**
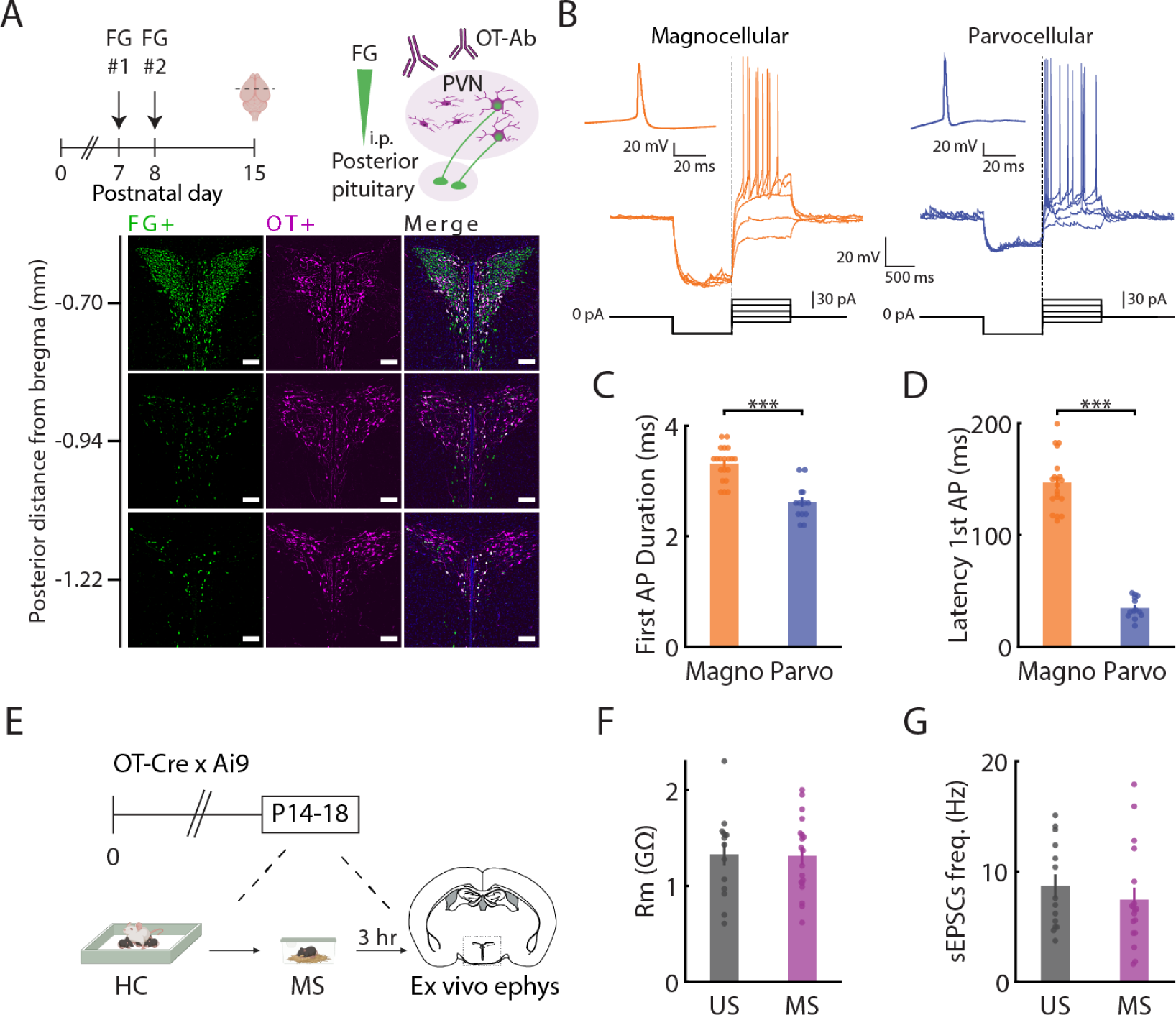
Anatomical and electrophysiological characteristics of OT neurons in the PVN following acute MS. **(A)** Anatomical identification of magno-OT and parvo-OT neurons in P15 mice, combining systemic FluoroGold infusion and OT immunoreactivity in the PVN (top). Bottom: Representative images of pup’s PVN in rostral-caudal axis, showing FG-labeled neurons (left; green) OT-expressing neurons (middle; magenta), and merged fluorescence (right). Magno-OT neurons (OT+/FG+ cells; white), parvo-OT neurons (OT+/FG-cells; magenta). Scale bar: 100 *µm*. **(B)** Representative traces of magnocelular (orange; magno-OT) and parvocellular (blue; parvo-OT) OT neurons in pups. (C) Latency to first AP for magno-OT (n = 20) and parvo-OT (n = 13) neurons. Student’s t-test *t*_(26.437)_ = 18.607, *P* = 1.03 *×* 10*^−^*^16^. (D) Duration of first AP for magno- and parvo-OT neurons. Student’s t-test *t*_(31)_ = 6.231, *P* = 6.36 *×* 10*^−^*^7^. (E) Schematics of *ex vivo* electrophysiology after MS. (F) Membrane resistance in OT neurons for MS (magenta) and US (gray) pups. Student’s t-test *t*_(31)_ = 0.103, *P* = 0.919. **(G)** Frequency of spontaneous excitatory postsynaptic currents (sEPSCs) in OT neurons for MS and US pups. Student’s t-test *t*_(30)_ = 0.796, *P* = 0.432. Data are presented as mean *±* s.e.m. (error bars). \*\*\**P <* 0.001.

**Extended Data Figure 4:**
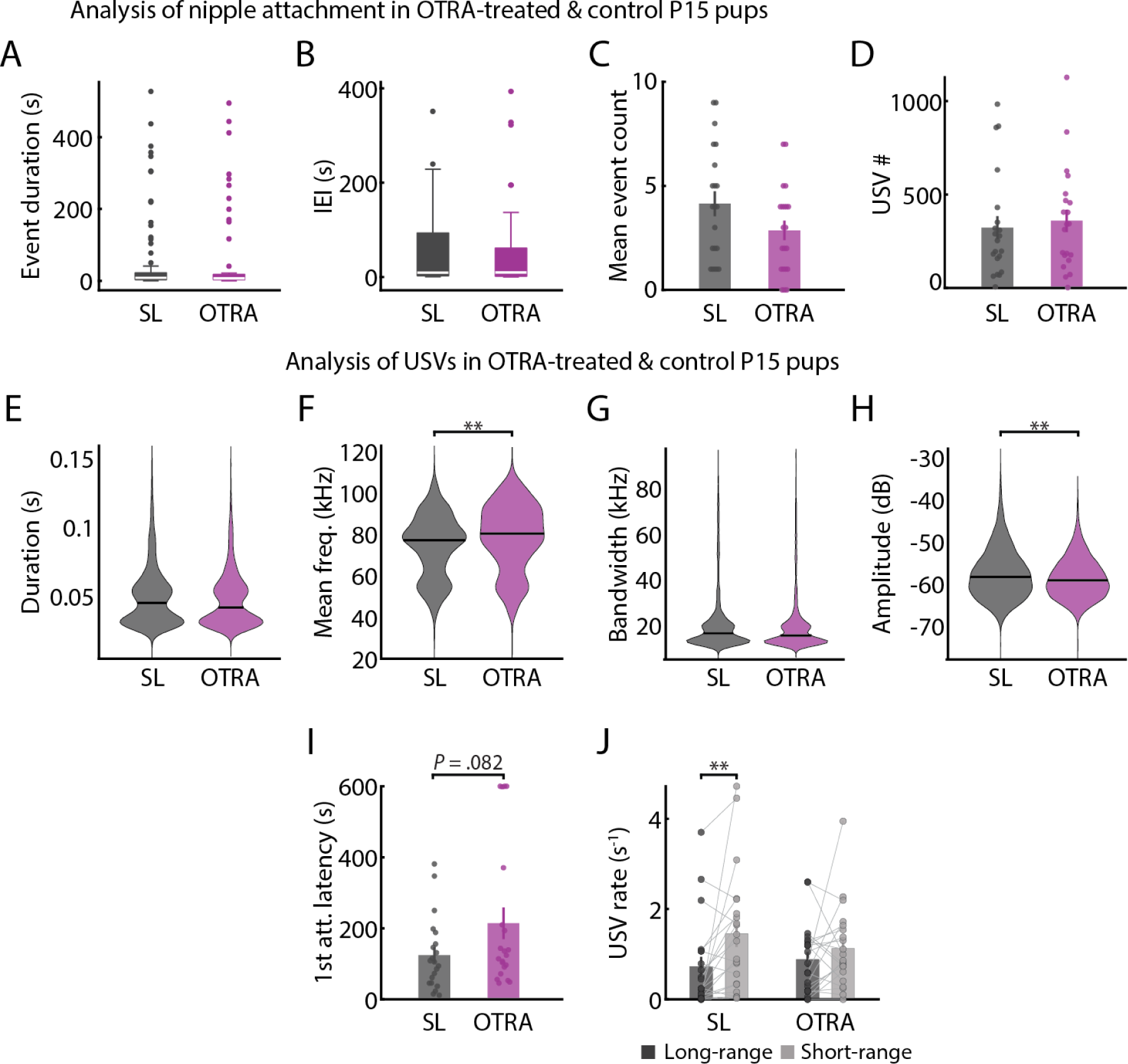
Analysis of nipple attachment and USV emission in OTRA-treated and control pups. **(A–B)** Distribution of (A) attachment event duration and (B) inter-event interval (IEI) for OTRA-treated (n*_OT RA_* = 21; magenta) and vehicle-treated (”SL”; n*_SL_* = 21; gray) pups. Box plot marks IQR (box edges) and median (white line), whiskers mark 1.5*±* IQR, dots mark outliers (¿1.5 IQR). Fitting a LME model for quantifying the differences in the distribution of events for OTRA and SL pups. (A) *t*_(145)_ = *−*0.764, *P* = 0.446; (B) *t*_(106)_ = 0.222, *P* = 0.824 **(C)** Number of attachment events during reunion, averaged across mice for OTRA and SL pups. Student’s t-test *t*_(40)_ = 1.704, 0.096. **(D)** Number of USVs emitted during reunion, averaged across mice for OTRA and SL pups. Student’s t-test *t*_(40)_ = *−*0.427, *P* = 0.671. **Extended Data Figure 4: (E–H)** Violin plots showing the properties of individual USVs emitted by OTRA and SL pups, characterized by (E) duration, (F) mean frequency, (G) bandwidth and (H) amplitude. Difference between distributions was calculated by fitting an LME model: *t*_(14317)_ = *−*1.44, *P* = 0.15; (F) *t*_(14317)_ = 2.641, *P* = 0.008; (G) *t*_(14317)_ = *−*1.251, *P* = 0.211; (H) *t*_(14317)_ = *−*2.779, *P* = 0.005. **(I)** Latency to first nipple attachment for OTRA and SL pups. Student’s t-test *t*_(29.358)_ = *−*1.8, *P* = 0.082. **(J)** Rate of long- and short-range USVs (dark and light gray, respectively) emitted before the first nipple attachment. Bonferroni-corrected two-sided Wilcoxon signed-rank test, SL: *W* = 34, *P* = 0.009; OTRA: *W* = 89, *P* = 0.714. Data in C,D,I,J are presented as mean *±* s.e.m. (error bars), \*\**P <* 0.01.

**Extended Data Figure 5:**
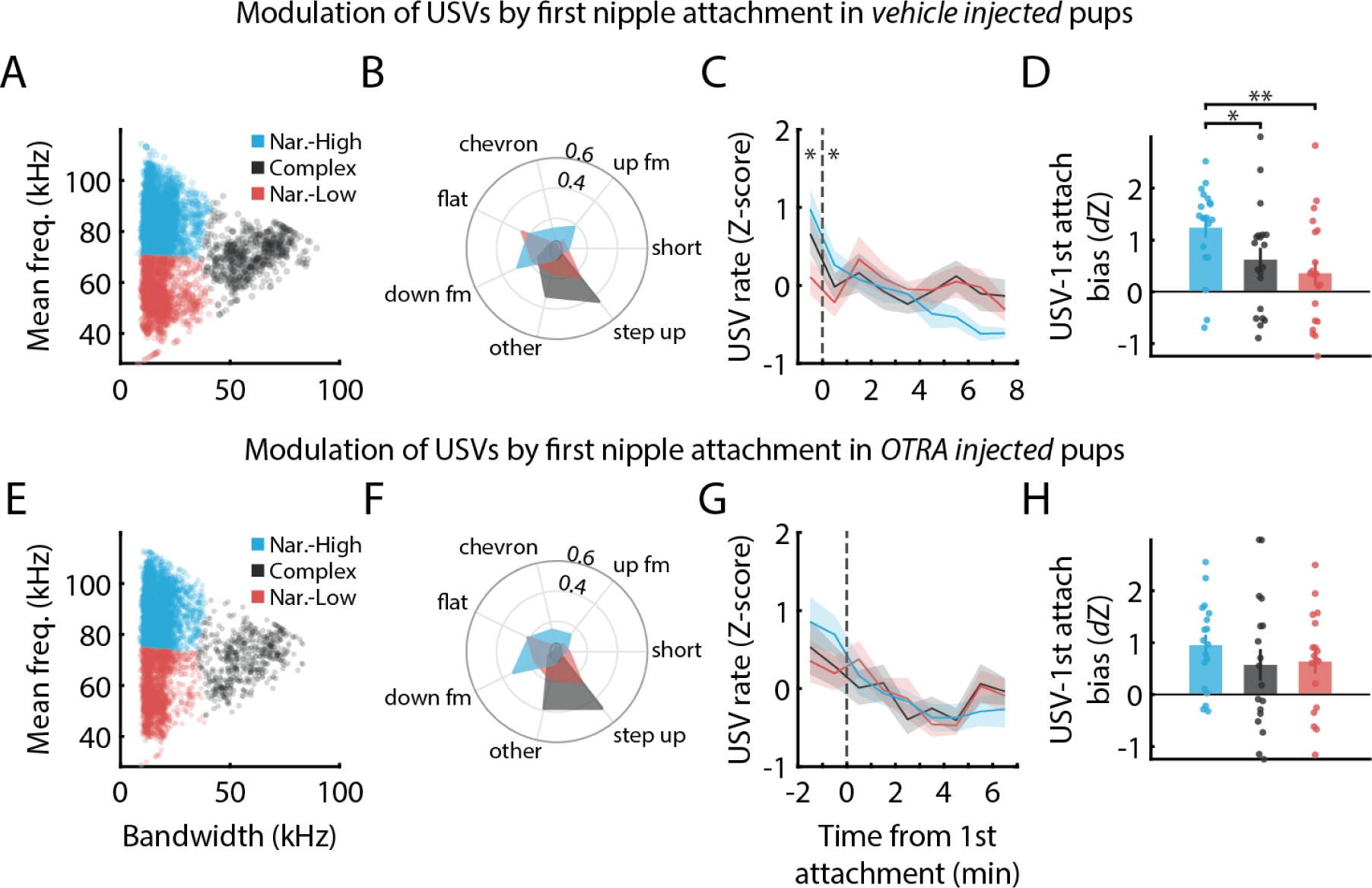
Blocking OT receptor during maternal separation attenuates attachment-associated USV modulation. **(A,E)** Individual USVs grouped into three clusters based on their bandwidth and mean frequency for control (A) and OTRA (E) pups, similar to the analysis in Fig. 1J. Colors in subsequent panels correspond with this cluster analysis. **(B,F)** Distribution of predefined vocal categories in each cluster for control (B) and OTRA (F) pups. **(C,G)** Normalized cluster-specific frequency of USV emission, aligned to first nipple attachment for control (C) and OTRA (G) pups. Two-way RM ANOVA examining the effect of cluster and time from first nipple attachment on USV emission with Bonferroni correction for post hoc comparisons. SL: *F_cluster_*_(2,38)_ = 3.178, *P_corrected_* = 0.106; *F_time_*_(4,76)_ = 3.124, P*_corrected_* = 0.039; F*_cluster×time_*_(8,152)_ = 3.238, P*_corrected_* = 0.004. OTRA: F*_cluster_*_(2,34)_ = 1.196, P*_corrected_* = 0.63; F*_time_*_(1,17)_ = 0.605, P*_corrected_* = 0.895; F*_cluster×time_*_(2,34)_ ^= 3^.^766,^ *P_corrected_* = 0.067. **(D,H)** Quantification of cluster-specific tendency to vocalize in relation to first nipple attachment for control (D) and OTRA (H) pups. RM ANOVA comparing clusters with Bonferroni-corrected post hoc comparisons: SL: *F_cluster_*_(2,38)_ = 9.299, *P_corrected_* = 0.001. OTRA: *F_cluster_*_(2,34)_ = 1.535, *P_corrected_* = 0.460. Data are presented as mean *±* s.e.m. (error bars and shaded area), \**P <* 0.05, \*\**P <* 0.01 For detailed statistical information, see Supplementary Table 1 and Supplementary Table 2.

**Extended Data Figure 6:**
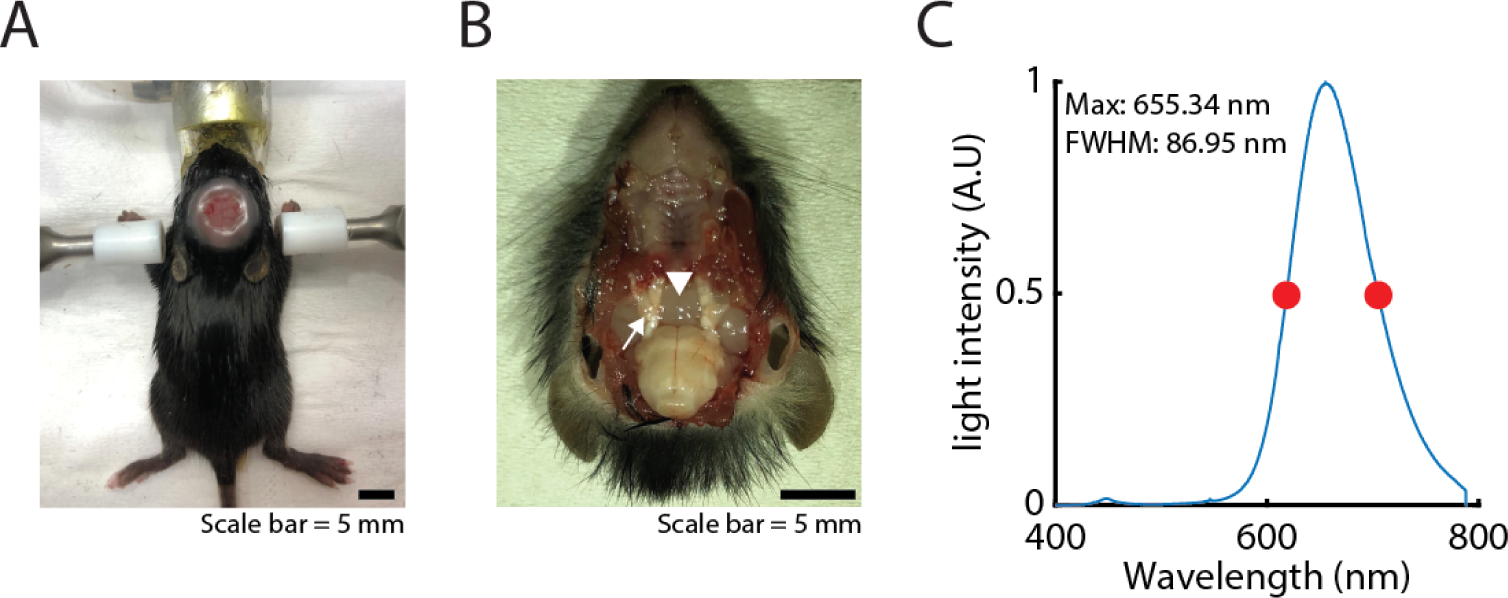
Intact-skull implant and illumination setup for transcranial optogenetics in postnatal mice. **(A)** Image depicts a P13 mouse with an intact-skull implant. **(B)** Dissection of the ventral hypothalamus (arrowhead), where a light sensor was placed for light measurements, as illustrated in Fig. 4K. Arrow: trigeminal ganglion. **(C)** Power spectral density (PSD) plots of the red LED arrays used in our transcranial optogenetic experiments, indicating the system’s max wavelength and full width at half maximum (FWHM; red dots).

**Extended Data Figure 7:**
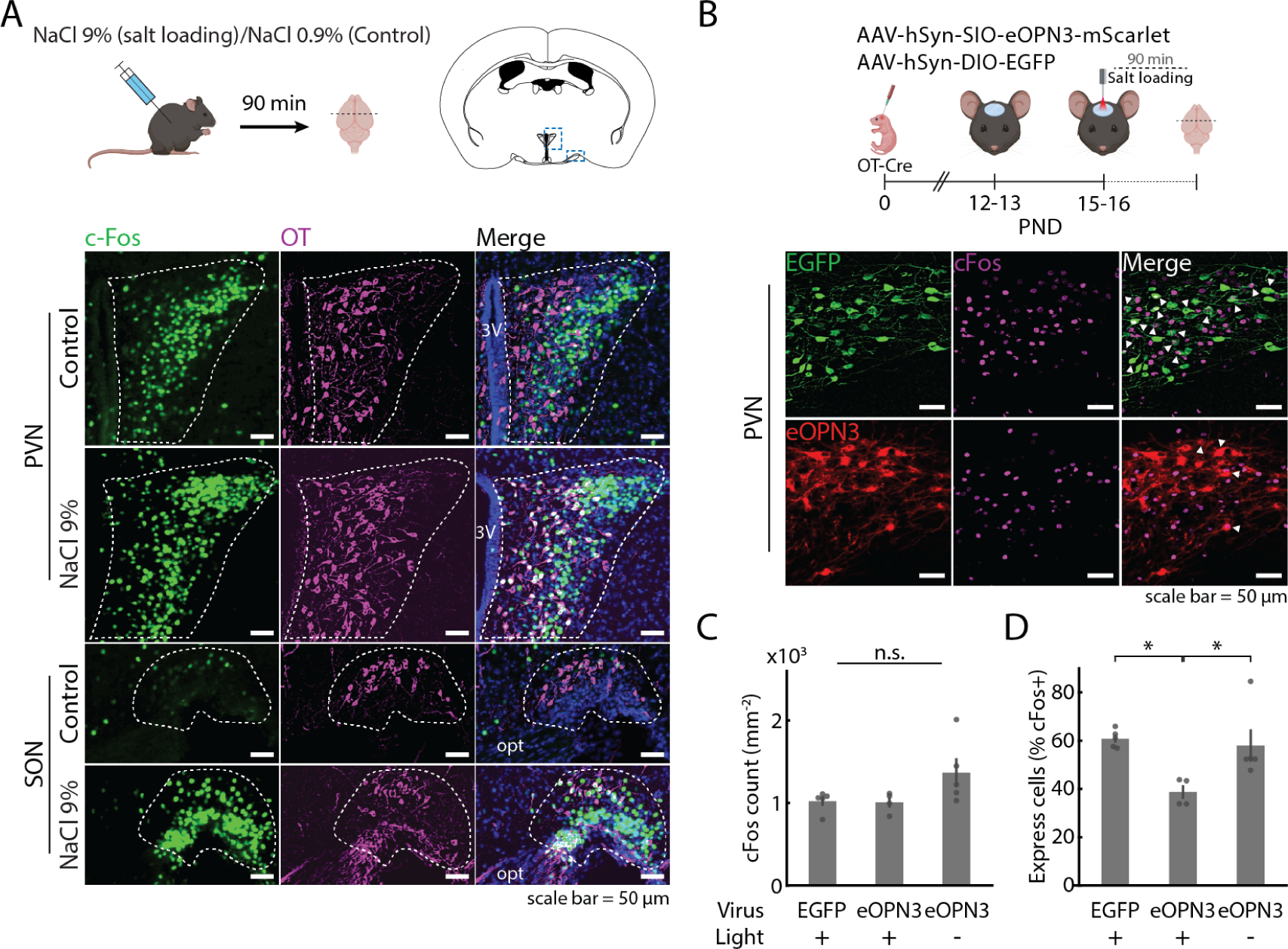
eOPN3 suppresses c-Fos expression in PVN-OT neurons after acute hyper-osmotic challenge in pups. **(A)** Top: Experimental paradigm for acute salt loading at P15. Pups were analyzed for cFos/OT immunoreactivty in sections of the PVN and SON (dashed line). Bottom: Representative images of control and salt-loaded pups, showing the overlap (white nuclei) of c-Fos (green) and OT (magenta) immunoreactivity in the PVN and SON. **(B)** Top: Schematic representation of the experimental approach for eOPN3-mediated transcranial silencing of OT neurons in anesthetized pups. Bottom: representative images of the PVN from OT-EGFP (green) and OT-eOPN3 (red) mice co-stained for c-Fos (magenta). Arrowheads indicate the overlap of c-Fos-positive neurons and EGFP- or eOPN3-positive neurons. **(C)** Quantification of c-Fos positive cells in the PVN for OT-EGFP (n = 5) and OT-eOPN3 (n = 4) pups after red light illumination (light +). A separate control of non-anesthetized eOPN3-expressing pups (n = 5) underwent salt loading without red light illumination (light -). One-way ANOVA *F*_(2,11)_ = 3.021, *P* = 0.09. **(D)** Percentage of EGFP-and eOPN3- expressing neurons positive for c-Fos immunoreactivity in the PVN. One-way ANOVA with post-hoc Bonferroni-corrected comparisons *F*_(2,11)_ = 6.489, *P* = 0.013. Data are presented as mean *±* s.e.m. (error bars), \**P <* 0.05. For detailed statistical information, see Supplementary Table 2.

**Extended Data Figure 8:**
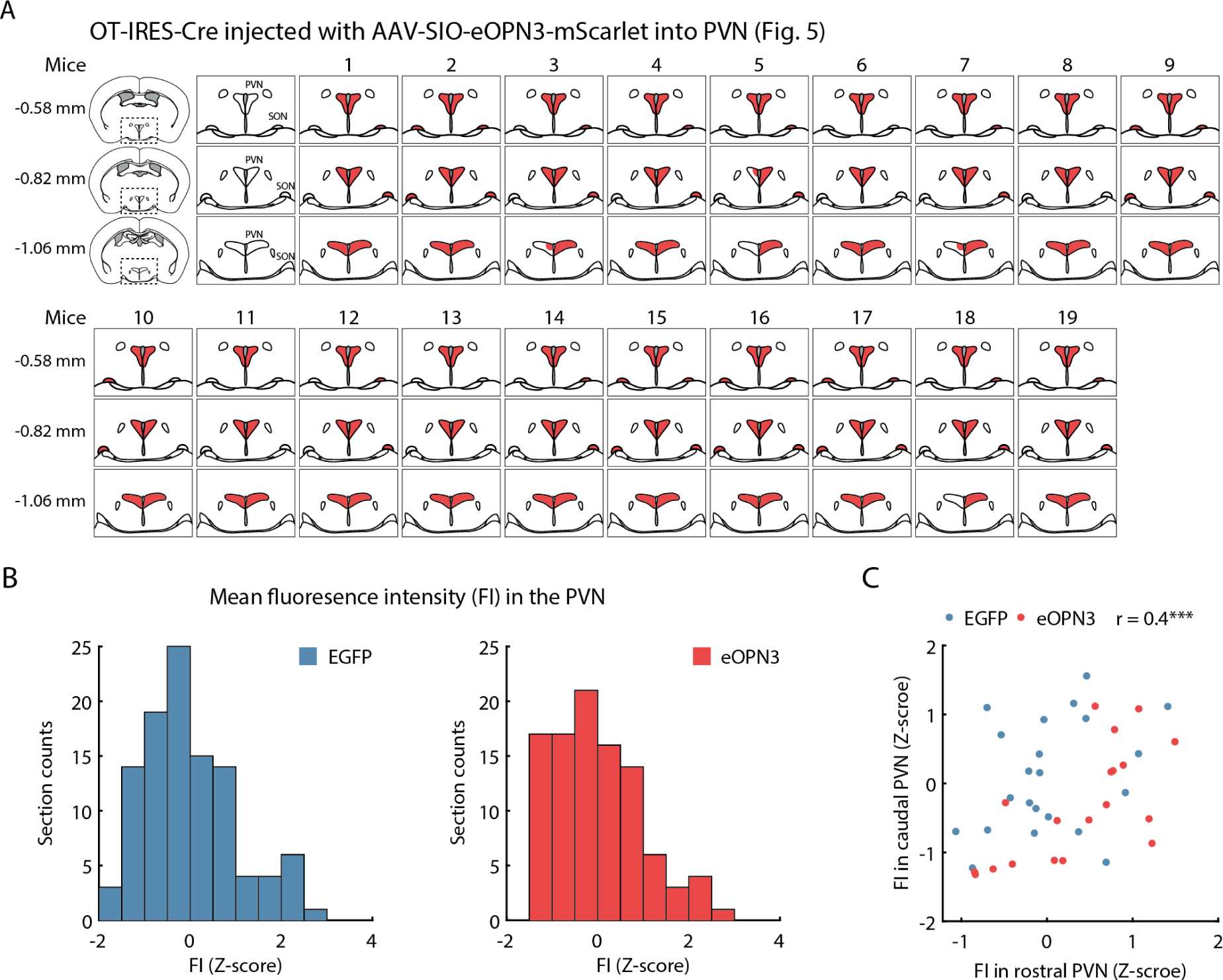
Histological analysis of viral expression of eOPN3-mScarlet and EGFP in the PVN of OT-IRES-Cre pups. **(A)** Schematics illustrating eOPN3-mScarlet expression (red shade) in the PVN and SON of OT-IRES-Cre pups for each animal in Fig. 5. Schematics corresponds with The Mouse Brain in Stereotaxic Coordinates Second Edition (Paxinos and Franklin). **(B)** Distribution of normalized mean fluorescence intensity in brain sections of the PVN for EGFP (blue; left; n = 105, N = 22) and eOPN3 (red; right; n = 99, N = 19). **(C)** Scatter plot of per mouse normalized mean fluorescence intensity of rostral and caudal PVN for EGFP (blue) and eOPN3 (red) pups. Pearson correlation coefficient calculated for all pups *r* = 0.4; *P* = 0.009

**Extended Data Figure 9:**
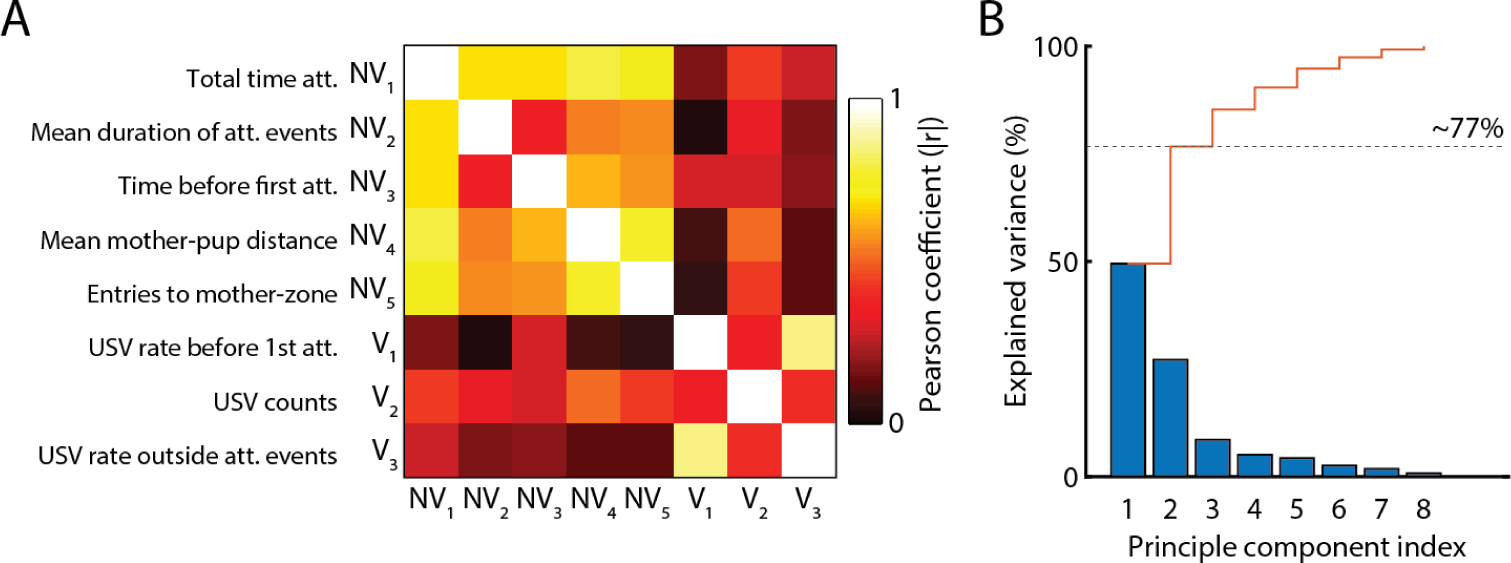
Principal component analysis of the vocal and non-vocal behavior of pups during reunion. **(A)** Correlation matrix of pups non-vocal (NV1-NV5) and vocal (V1-V3) behaviors during reunion with the dam for eOPN3 and EGFP pups, related to experiment in Fig. 5. Color map indicates the absolute Pearson correlation coefficient. **(B)** Explained variance by principal components (PC1-PC8) derived from vocal and non-vocal behaviors described in **A**, overlaid with cumulative explained variance (red line).

**Table 1:** Classification of USVs - USVs emitted during reunion were classified into three clusters based on their bandwidth (bw) and mean frequency (f).

| Figure | Group | Narrow-High<sup>1</sup> |  | Complex<sup>2</sup> |  | Narrow-Low<sup>3</sup> |  |
| --- | --- | --- | --- | --- | --- | --- | --- |
|  |  | bw | f | bw | f | bw | f |
| Fig. 1J | MS | n = 1388 |  | n = 221 |  | n = 1020 |  |
| | | $\bar{x} = 16.73$ | $\bar{x} = 87.64$ | $\bar{x} = 58.14$ | $\bar{x} = 72.23$ | $\bar{x} = 16.02$ | $\bar{x} = 55.93$ |
| | | $S = 0.14$ | $S = 0.22$ | $S = 0.75$ | $S = 0.44$ | $S = 0.15$ | $S = 0.2$ |
| Ext. Data Fig. 5A | SL | n = 4110 |  | n = 607 |  | n = 2056 |  |
| | | $\bar{x} = 17.28$ | $\bar{x} = 84.1$ | $\bar{x} = 59.54$ | $\bar{x} = 70.74$ | $\bar{x} = 17.05$ | $\bar{x} = 55.67$ |
| | | $S = 0.09$ | $S = 0.13$ | $S = 0.46$ | $S = 0.29$ | $S = 0.13$ | $S = 0.16$ |
| Ext. Data Fig. 5E | OTRA | n = 4709 |  | n = 573 |  | n = 2264 |  |
| | | $\bar{x} = 16.96$ | $\bar{x} = 88.62$ | $\bar{x} = 56.58$ | $\bar{x} = 70.67$ | $\bar{x} = 16.06$ | $\bar{x} = 59.18$ |
| | | $S = 0.08$ | $S = 0.13$ | $S = 0.47$ | $S = 0.35$ | $S = 0.1$ | $S = 0.2$ |

<sup>1</sup> USVs with narrow bandwidth and high mean frequency.
<sup>2</sup> USVs with varied bandwidth and medium mean frequency.
<sup>3</sup> USVs with narrow bandwidth and low mean frequency.

**Table 2:** Comprehensive statistical information of all post-hoc comparisons presented in this publication.

| Figure | Test | Comparison | P-value |
| --- | --- | --- | --- |
| Fig. 1K | Bonferroni corrected multiple RM ANOVA tests | chevron | 0.030 |
|  |  | down fm | 0.014 |
| | | flat | $5.9 \times 10^{-6}$ |
|  |  | short | 0.004 |
| | | step up | $4.4 \times 10^{-4}$ |
|  |  | up fm | 0.012 |
|  |  | other | 0.078 |
| Fig. 1L | Bonferroni corrected post-hoc comparisons<br>(following RM ANOVA) | Narrow-High & Complex | 0.074 |
|  |  | Narrow-High & Narrow-Low | 0.004 |
|  |  | Complex & Narrow-Low | 1.000 |
| Fig. 1M | Bonferroni corrected post-hoc comparisons between<br>clusters for valid time bins<br>(following two-way RM ANOVA) | [-60:0] sec | 0.009 |
|  |  | [0:60] sec | 1.000 |
|  |  | [60:120] sec | 1.000 |
|  |  | [120:180] sec | 1.000 |
|  |  | [180:240] sec | 1.000 |
|  |  | [240:300] sec | 0.121 |
|  |  | [300:360] sec | 1.000 |
|  |  | [360:420] sec | 0.047 |
| Fig. 1N | Bonferroni corrected post-hoc comparisons<br>(following RM ANOVA) | Narrow-High & Complex | 0.044 |
|  |  | Narrow-High & Narrow-Low | 0.028 |
|  |  | Complex & Narrow-Low | 1.000 |

Table 2 – continued from previous page
| Figure | Test | Comparison | P-value |
| --- | --- | --- | --- |
| Fig. 2B | Bonferroni corrected post-hoc comparisons<br>(following one-way ANOVA in PVN) | MS & US | 0.039 |
|  |  | MS & Reunion | 0.018 |
|  |  | US & Reunion | 1.000 |
|  | Bonferroni corrected post-hoc comparisons<br>(following one-way ANOVA in SON) | MS & US | 0.021 |
|  |  | MS & Reunion | 0.016 |
|  |  | US & Reunion | 1.000 |
| Fig. 3C | Bonferroni corrected post-hoc comparisons<br>(following mixed-design RM ANOVA) | SL: No attach & Attach | 0.024 |
|  |  | OTRA: No attach & Attach | 1.000 |
| Fig. 3G | Bonferroni corrected post-hoc comparisons<br>(following RM ANOVA) | Narrow-High & Complex | 0.989 |
|  |  | Narrow-High & Narrow-Low | 0.003 |
|  |  | Complex & Narrow-Low | 0.273 |
| Fig. 5C | Bonferroni corrected post-hoc comparisons<br>(following mixed-design RM ANOVA)<br>$F_{sex(1,37)} = 0.37, P = 0.547; F_{group \times sex(1,37)} = 1.79;$<br>$P = 0.189; F_{sex \times phase(1,37)} = 0.336, P = 0.566$ | EGFP: 1<sup>st</sup> half & 2<sup>nd</sup> half | 0.001 |
|  |  | eOPN3: 1<sup>st</sup> half & 2<sup>nd</sup> half | 0.865 |
| Fig. 5F | General linear model (GLM)<br>$F_{(5,31)} = 0.323, P = 0.895$<br>$R^2 = 0.05, R^2_{adj.} = -0.103$ | Fluorescence intensity<br>$t_{31} = -0.172$ | 0.864 |
| | | MS USV index<br>$t_{31} = 0.096$ | 0.924 |
| | | Group<br>$t_{31} = -0.162$ | 0.873 |
| | | Sex<br>$t_{31} = -0.531$ | 0.599 |
| | | Group X Sex<br>$t_{31} = -0.309$ | 0.759 |

Table 2 – continued from previous page
| Figure | Test | Comparison | P-value |
| --- | --- | --- | --- |
| Fig. 5G | General linear model (GLM)<br>$F_{(5,31)} = 3.958, P = 0.007$<br>$R^2 = 0.39, R^2_{adj.} = 0.291$ | Fluorescence intensity<br>$t_{31} = 0.924$ | 0.363 |
| | | MS USV index<br>$t_{31} = 2.59$ | 0.015 |
| | | Group<br>$t_{31} = -2.657$ | 0.012 |
| | | Sex<br>$t_{31} = -3.405$ | 0.002 |
| | | Group X Sex<br>$t_{31} = 3.051$ | 0.005 |
|  | Bonferroni corrected post-hoc comparisons of GLM | EGFP: F & M | 0.007 |
|  |  | eOPN3: F & M | 1.000 |
|  |  | F: EGFP & eOPN3 | 0.049 |
|  |  | M: EGFP & eOPN3 | 0.485 |
| Fig. 5I | General linear model (GLM)<br>$F_{(5,31)} = 0.427, P = 0.826$<br>$R^2 = 0.064, R^2_{adj.} = -0.086$ | Fluorescence intensity<br>$t_{31} = -0.002$ | 0.998 |
| | | MS USV index<br>$t_{31} = -0.296$ | 0.769 |
| | | Group<br>$t_{31} = 0.212$ | 0.834 |
| | | Sex<br>$t_{31} = -0.019$ | 0.985 |
| | | Group X Sex<br>$t_{31} = -0.935$ | 0.357 |

Table 2 – continued from previous page
| Figure | Test | Comparison | P-value |
| --- | --- | --- | --- |
| Fig. 5J | General linear model (GLM)<br>$F_{(5,31)} = 2.892, P = 0.03$<br>$R^2 = 0.318, R^2_{adj.} = 0.208$ | Fluorescence intensity<br>$t_{31} = 1.158$ | 0.256 |
| | | MS USV index<br>$t_{31} = 1.843$ | 0.075 |
| | | Group<br>$t_{31} = -2.37$ | 0.024 |
| | | Sex<br>$t_{31} = -2.914$ | 0.007 |
| | | Group X Sex<br>$t_{31} = 2.1$ | 0.044 |
|  | Bonferroni corrected post-hoc comparisons of GLM | F: EGFP & eOPN3 | 0.048 |
|  |  | M: EGFP & eOPN3 | 1.000 |
| Ext. Data<br>Fig. 5C | Bonferroni corrected post-hoc comparisons between clusters for valid time bins (following two-way RM ANOVA) | SL: [-60:0] sec | 0.023 |
|  |  | SL: [0:60] sec | 0.042 |
|  |  | SL: [60:120] sec | 1.000 |
|  |  | SL: [120:180] sec | 1.000 |
|  |  | SL: [180:240] sec | 1.000 |
| Ext. Data<br>Fig. 5D | Bonferroni corrected post-hoc comparisons (following RM ANOVA) | SL: Narrow-High & Complex | 0.028 |
|  |  | SL: Narrow-High & Narrow-Low | 0.003 |
|  |  | SL: Complex & Narrow-Low | 0.501 |
| Ext. Data<br>Fig. 7D | Bonferroni corrected post-hoc comparisons (following one-way ANOVA) | EGFP-on & eOPN3-on | 0.019 |
|  |  | EGFP-on & eOPN3-off | 1.000 |
|  |  | eOPN3-on & eOPN3-off | 0.041 |

